# Conserved Fibroblast Subsets Drive Shared Fibrotic Mechanisms in Osteoarthritis Synovium and Systemic Sclerosis Skin

**DOI:** 10.64898/2026.10.05.756653

**Authors:** Aanya Mohan, Michael D. Newton, Neil Bhate, Alexandra Khmelevskaya, Tareq M. Hanna, Edward DiCarlo, Caroline Ospelt, Dana Orange, John Varga, Tristan Maerz

**Affiliations:** Institute for Biomechanics, Department of Health Sciences and Technology, ETH Zürich, Zürich, Switzerland; Department of Orthopaedic Surgery, University of Michigan, Ann Arbor, MI, USA; Department of Rheumatology, University Hospital Zürich, University of Zürich, Zürich, Switzerland; Wayne State University School of Medicine, MI, USA; Hospital for Special Surgery, New York, NY, USA; Rockefeller University, New York, NY, USA; Department of Biomedical Engineering, University of Michigan, Ann Arbor, MI, USA; Division of Rheumatology, Department of Internal Medicine, University of Michigan, Ann Arbor, MI, USA

## Abstract

Osteoarthritis (OA) and systemic sclerosis (SSc) are diseases with distinct etiologies but similar disease phenotypes, most prominently fibrosis, a process governed by the pathological activation of fibroblasts. Mechanisms underpinning fibrosis in these diseases are widely studied, though have yet to be directly compared. We employed single-cell RNA-seq and high-resolution spatial transcriptomics to describe the functional plasticity and the range of fibroblast activation states in both diseases. We identified disease-enriched fibroblast subsets conserved in both diseases, including fibrogenic myofibroblast-like cells, chemokine- and cytokine-expressing inflammatory fibroblasts, and lining-like fibroblasts. These subsets exhibited convergent fibrotic programs also observed in idiopathic pulmonary fibrosis, as well as overlapping osteochondral transcriptional programs. Xenium 5K spatial transcriptomics revealed that conserved, disease-associated fibroblasts reside in highly similar anatomic niches in OA synovium and SSc skin. These consisted of a fibrotic ECM-rich niche harboring myofibroblast-like cells and an inflammatory niche containing inflammatory fibroblasts adjacent to infiltrating immune cells. Disease-associated transcriptional remodeling converged on shared functional pathways and common putative upstream molecular regulators. These findings identify conserved cellular states, spatial niches, and regulatory programs across OA and SSc, suggesting that fibrotic disease is underpinned by shared pathogenic stromal states and candidate targets for therapeutic strategies are applicable across both diseases.

## Introduction

Fibrosis is a pathological feature common to multiple rheumatic and musculoskeletal diseases, is strongly associated with morbidity and mortality, and represents a substantial global healthcare burden^1^. Two disparate conditions in which fibrosis contributes substantially to disease pathology are osteoarthritis (OA) and systemic sclerosis (SSc). OA is a complex whole-joint disease characterized by articular destruction, joint pain, and pathological remodeling, and inflammation of the synovium is increasingly understood as a key contributor to OA pathogenesis. SSc is a systemic autoimmune disease characterized by excessive collagen deposition across several organs including the skin. Despite differences in anatomical location and disease presentation, OA synovium and SSc skin display several convergent pathological features including excessive extracellular matrix (ECM) accumulation and focal inflammatory infiltrates (**Fig.-1A-B**), vascular remodeling, and ectopic mineralization^2–5^. Furthermore, patients with SSc have been shown to exhibit evidence of synovitis and pain in synovial joints^6,7^. These observations raise the possibility that common stromal mechanisms underlie pathological tissue remodeling across both conditions.

**Figure 1:**
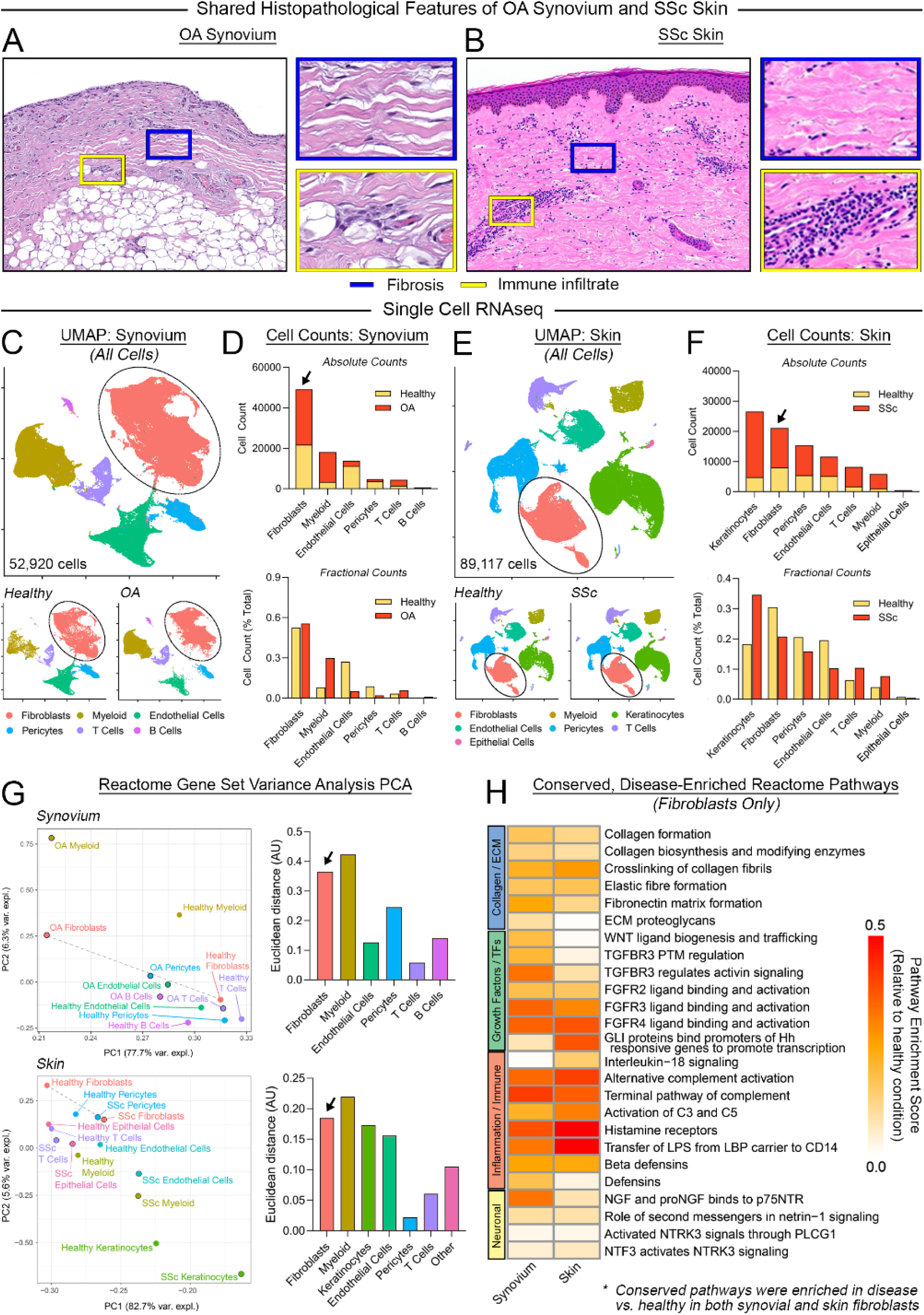
Fibroblasts in synovium and skin exhibit a high degree of plasticity. **(A,B)** Representative images of H&E-stained histological sections of OA synovium (left, courtesy of Edward DiCarlo, Hospital for Special Surgery) and SSc skin (right, courtesy of May Chan, University of Michigan) illustrating convergent histopathological features of OA and SSc, including areas of fibrotic tissue deposition (blue box) and focal immune infiltrate (yellow box). **(C-D)** Clustering and annotation (C) and counts/proportions in healthy and OA tissue (D) of major cell types in scRNA-seq data of synovium (n=52,920 total cells from n=5 datasets) **(E-F)** Clustering and annotation (E) and counts/proportions in healthy and SSc tissue (F) of major cell types in scRNA-seq data of skin (n=89,117 total cells from n=4 datasets) **(G)** Principal component analysis (PCA) of Reactome pathway enrichment profiles for major cell types in healthy and diseased synovium (top) and skin (bottom). Left: Points on PCA plots represent cell type and healthy/disease condition, and dashed lines connect matched healthy and diseased cell types. Right: Euclidean distances between matched healthy and diseased cell types quantify the magnitude of disease-associated functional divergence. **(H)** Heatmap of Reactome pathways positively enriched in both OA synovium and SSc skin fibroblasts relative to respective healthy conditions. Rows indicate conserved disease-enriched pathways, grouped by thematic biological functions.

Fibrosis is characterized by the aberrant overproduction of collagen and other ECM molecules, progressively disrupting normal tissue architecture and homeostatic function. Fibroblasts are the primary cell type responsible for fibrotic ECM production across various tissues and conditions^8^. Conventionally viewed as unitary matrix-producing and -remodeling cells, fibroblasts are increasingly recognized as critical regulators of immune, endothelial, and other stromal cells via dynamic crosstalk^9^. Their role as disease drivers, rather than responders to altered tissue homeostasis, is under active investigation across rheumatic and musculoskeletal conditions including OA and SSc^8–19^. Single-cell RNA sequencing (scRNA-seq) has revolutionized the understanding of fibroblast identity and heterogeneity in health and disease - fibroblasts are now recognized as highly plastic cells existing on a multidimensional spectrum of phenotypic states, dictated by anatomic location and disease status^20,21^. Extending the insights gained from scRNA-seq, spatial transcriptomics (particularly technologies enabling true single-cell resolution) have the potential to further bridge the gap between cellular mechanisms and disease phenotypes, defining disease-associated niches across tissues and disease states. Leveraging these technologies to uncover shared molecular mechanisms and spatial niches governing fibroblast activation in fibrotic diseases may inform novel multi-disease therapeutic strategies.

Currently, no disease-modifying treatments exist for OA or SSc, emphasizing the need to further understand the cellular and molecular mechanisms underpinning patient outcome-linked disease manifestations such as progressive fibrosis and chronic inflammation. Given the phenotypic similarities between OA and SSc, we postulated that conserved stromal cell subsets orchestrate similar pro-fibrotic programs in both diseases. Prior studies have demonstrated similar fibroblast subsets, and core elements of pro-fibrotic fibroblast activation, shared across diverse tissues and disease contexts including cancer, lung, and kidney fibrosis^10,11,16,17,20^. While these studies have identified common signaling pathways associated with fibroblast activation, including TGFβ, WNT, and FGF signaling, none have directly compared OA synovium with SSc skin.

To this end, we generated an integrated single-cell atlas of healthy and diseased synovium and skin encompassing 69,047 human fibroblasts. We characterized multiple transcriptionally-distinct fibroblast populations shared between skin and synovium to determine which subsets underwent disease-associated expansion, fibrogenic signaling, and disease-induced changes in gene expression in OA and SSc. We leveraged Xenium 5K spatial transcriptomics to define the spatial niches of conserved, disease-associated fibroblast subsets. Finally, we identified convergent signaling programs and putative regulators associated with fibroblast activation states across OA and SSc. Our findings demonstrate previously unrecognized parallel fibrotic mechanisms, extending the knowledge of mechanisms of fibrosis across diverse diseases and informing new avenues regarding their treatment.

## Results

### Fibroblasts in synovium and skin exhibit a high degree of disease-associated plasticity

To characterize conserved cellular and molecular programs in OA and SSc, we integrated 9 publicly available scRNA-seq datasets comprising healthy and diseased synovium and skin samples (five synovium datasets, constituting n=11 healthy/non-OA and n=10 OA subjects^12,18,22–24^; four skin datasets, constituting n=15 healthy/non-SSc and n=38 SSc subjects^14,25–27^). OA synovium samples were taken during total knee replacement surgery, while healthy synovial samples were obtained postmortem, during ACL reconstruction, or during arthroscopic procedures in a non-OA knee joint; healthy and SSc skin samples were obtained as punch biopsies (**Supp.-Fig.-1A-B**). Within each dataset, stringent quality control was performed by sample to isolate high-quality cells, which were integrated and clustered to identify major cell types (**Fig.-1C-F, Supp.-Fig.- 1C-E**). Fibroblasts were identified as *PDGFRA+*, *PTPRC/CD45-* (**Supp.-Fig.-1F**) and constituted the most abundant cell type in synovium and the second most abundant cell type in skin in both healthy and diseased conditions (**Fig.-1D, F, Supp.-Fig.-1D**).

To broadly assess disease-associated functional diversification of major cell types, Reactome pathway analysis of within-cluster gene expression was performed, and results for enriched terms underwent principal component analysis (PCA) analysis (**Fig.-1G**). Calculating the *n*-dimensional Euclidean distance between healthy and diseased conditions across principal components thus provided a quantitative measure of overall disease-associated functional diversification. Fibroblasts exhibited a high degree of disease-associated plasticity in OA and SSc, second in magnitude to myeloid cells (**Fig.-1G**). Whereas high plasticity in myeloid cells is attributable to recruitment of systemically-derived cells into diseased tissue^28^, fibroblast plasticity represents expansion, activation, and diversification of tissue-resident cells. Compared to healthy fibroblasts, OA synovium and SSc skin fibroblasts were similarly enriched in Reactome terms related to fibrotic remodeling, including deposition and crosslinking of collagen and other ECM constituents, FGF and WNT pathway activation, and innate immune activation (complement pathway, beta-defensin expression) (**Fig.-1H**). These findings indicate a marked disruption of fibroblast homeostasis in both OA and SSc.

### OA synovium and SSc skin harbor conserved, disease-associated fibroblast subsets

To define specific fibroblast subpopulations underpinning the high degree of plasticity observed in OA and SSc, we extracted and integrated fibroblasts from both tissues and conditions, yielding a dataset comprised of 69,047 fibroblasts clustered into 12 transcriptionally distinct subsets based on cluster-specific marker genes (**Fig.-2A-B, Supp.-Fig.-2**). Relative abundance testing (*scProportionTest*, FDR < 0.05, |log2FD| > 0.58) defined clusters enriched in OA, SSc, or both, compared to healthy (**Fig.-2C**). Inferred pathway activity scores (**Fig.-2D**) and biological pathway activation (**Fig.-2E**) further characterized and functionally annotated fibroblast subsets.

**Figure 2:**
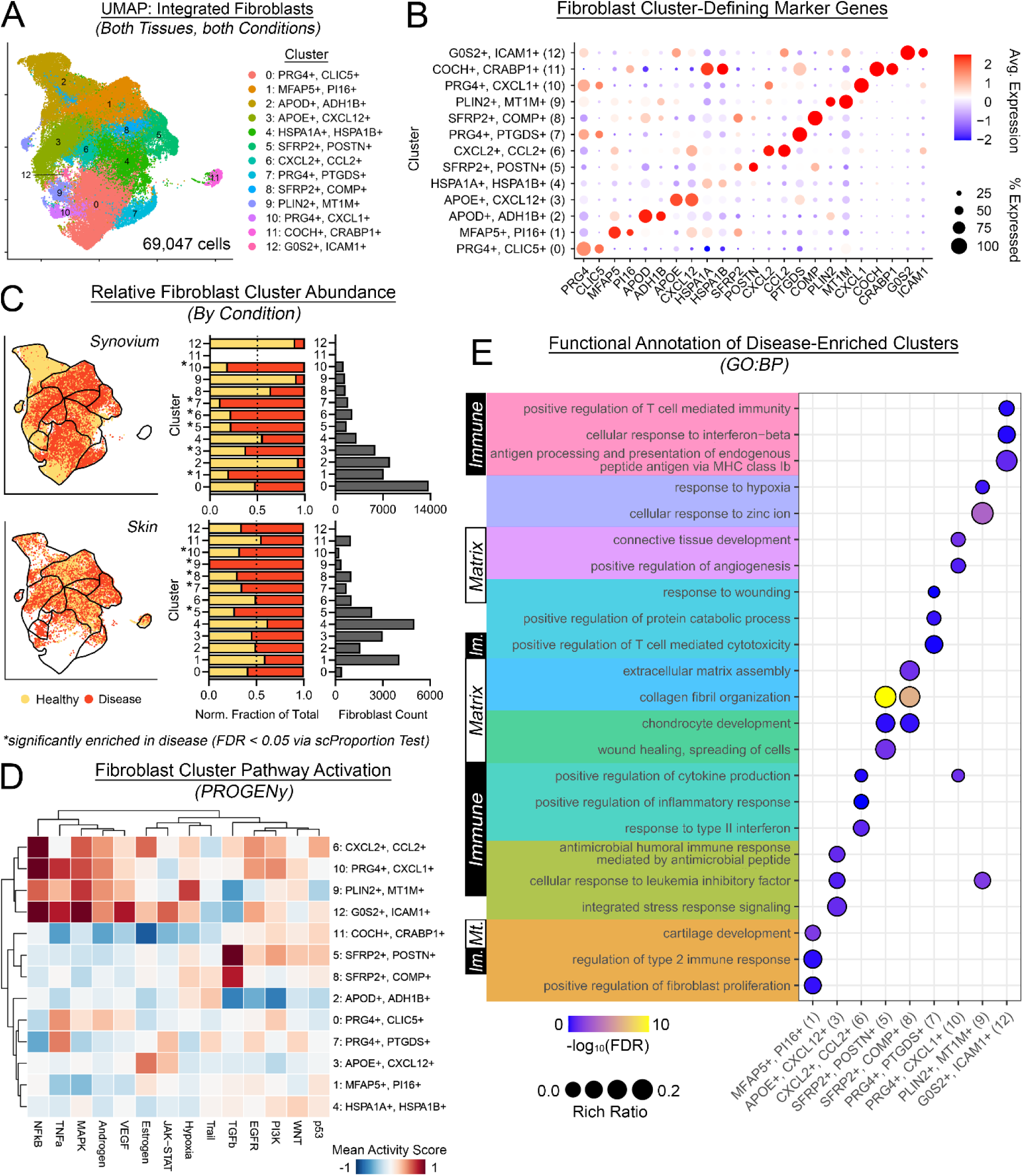
OA synovium and SSc skin harbor conserved, disease-associated fibroblast subsets. **(A)** UMAP of 69,047 human fibroblasts integrated across synovium and skin and across healthy and diseased conditions, identifying 13 transcriptionally distinct fibroblast clusters. Clusters are annotated by cluster-defining marker genes. **(B)** Dot plot of cluster-defining marker gene expression across fibroblast subsets. Dot color indicates scaled average expression, and dot size indicates the percentage of cells expressing the gene. **(C)** Relative abundance of fibroblast clusters in synovium (top) and skin (bottom), stratified by disease condition. Left: UMAPs illustrating the distribution of healthy and diseased cells. Right: Stacked bars of normalized fraction of cells from healthy and disease conditions across clusters, with adjacent bars showing absolute counts. Asterisks denote clusters significantly enriched in disease relative to the corresponding healthy tissue, as determined by *scProportionTest* using permutation testing and bootstrapping (FDR < 0.05, |log2FD| > 0.58). **(D)** Heatmap of inferred PROGENy pathway activity across fibroblast clusters. Values represent mean activity scores via a multivariate linear model; rows and columns are hierarchically clustered. **(E)** Gene Ontology-Biological Process (GO:BP) enrichment analysis of cluster-defining genes across disease-enriched fibroblast clusters, grouped by cluster and labeled with broad functional category. Dot size indicates rich ratio, and dot color indicates enrichment significance (−log10(FDR)) via PANTHER overrepresentation testing using positively enriched cluster-defining genes (*P_adj_* < 0.05; expression in >50% of cells within the cluster).

Certain fibroblast subsets were highly specific to synovium or skin (**Fig.-2C**). Cluster 0 (*PRG4+, CLIC5+*) was largely confined to synovium and expressed classical markers of synovial lining fibroblasts^13^ (**Fig.-2B**). Cluster 11 (*COCH+, CRABP+*) was found uniquely in skin and expressed classical markers of hair follicle-associated fibroblasts^11^. All other fibroblast clusters were meaningfully detected in synovium and skin, with each tissue contributing >20% of cells in each cluster after normalizing for cell number (**Supp.-Fig.-3F**).

Cluster 1 (*MFAP5+, PI16+*) expressed a gene signature consistent with previously-described *PI16+/CD34+/MFAP5+* pan-tissue progenitor or “universal” fibroblasts, shown to give rise to multiple stromal lineages^17,20^ (**Fig.-2E**). Cluster 1 was enriched in OA synovium compared to healthy synovium (**Fig.-2C**).

Cluster 5 (*SFRP2+, POSTN+*) and Cluster 8 (*SFRP2+, COMP+*) represent two related populations of ECM-enriched fibroblasts with predicted fibrogenic function. Cluster 5 was enriched in both OA synovium and SSc skin compared to healthy controls (**Fig.-2C, Supp.-Fig.-3F**) and exhibited enrichment of pathways including collagen fibril organization and wound healing (**Fig.-2E**), along with expression of classical gene markers for myofibroblasts including *SFRP2, POSTN, COMP, COL8A1, TNC, LOX, and TAGLN*^11,14,19^ (**Supp.-Fig.-3B**). TGFβ signaling, a canonical pro-fibrotic and myofibroblast-linked pathway^29^, was strongly activated in cluster 5 (**Fig.-2D**). Cluster 8 (*SFRP2+, COMP+*) was transcriptionally and functionally similar to fibrogenic cluster 5 (*SFRP2+, POSTN+*), enriched in pathways related to ECM assembly, exhibiting high expression of myofibroblast-related genes, and activation of TGFβ (**Fig.-2D-E, Supp.-Fig.-3B**). *SFRP2+ COMP+* skin fibroblasts arising from *PI16+* stromal progenitors (analogous to *MFAP5+, PI16+* fibroblasts in this data) were recently demonstrated to be enriched in regions of high matrix stiffness^19^. Cluster 8 (*SFRP2+, COMP+*) was enriched specifically in SSc skin (**Fig.-2C, Supp.-Fig.-3F**). Despite shared expression of a myofibroblast-like signature, clusters 5 and 8 demonstrated unique expression profiles (**Supp.-Fig.-3D**), corroborating recent studies distinct populations of fibrogenic, myofibroblast-like cells in RA synovium and SSc skin^6,10,17,30^.

Clusters 3 (*APOE+, CXCL12+*) and 6 (*CXCL2+, CCL2+*) were identified as immune-signaling fibroblasts, exhibiting cytokine and chemokine gene expression and enrichment of several immune signaling-related pathways (**Fig.-2D-E, Supp.-Fig.-3C**). Cluster 3 was enriched for pathways related to adaptive immune system activation, while cluster 6 was enriched for pathways relating to general inflammatory response and type II interferon signaling. Based on pathway activity analyses, cluster 6 was enriched for NFκB and MAPK signaling whereas cluster 3 was enriched for JAK-STAT and estrogen pathway signaling (**Fig.-2D**), suggesting distinct signaling pathways may underlie immune signaling within these clusters. Clusters 3 and 6 were specifically enriched in OA synovium compared to healthy synovium (**Fig.-2C**).

Clusters 7 (*PRG4+, PTGDS+*) and 10 (*PRG4+, CXCL1+*) represented further lining-like/lining-associated fibroblasts and were enriched in both OA synovium and SSc skin compared to healthy controls (**Fig.-2C, Supp.-Fig.-3F**). In the synovium, clusters 7 and 10 strongly expressed *PRG4* (encoding the joint-lubricating protein lubricin) and *CLIC5*, consistent with synovial lining fibroblasts^13^. These clusters were also meaningfully present in skin, expressing most cluster-defining genes, though notably not *PRG4* or *CLIC5* (**Supp.-Fig.-2G, 3E**). Cluster 7 was enriched for pathways related to both wound response and immune signaling, while cluster 10 was enriched for terms related to connective tissue development and angiogenesis. Both clusters exhibited high predicted TNFα pathway activity, a pro-inflammatory signaling pathway with established roles in OA and SSc^7,31^ (**Fig.-2D**).

The remaining smaller populations exhibited specialized or tissue-biased states. Cluster 12 (*G0S2+, ICAM+*) exhibited low overall incidence in both tissues, precluding downstream analyses. Cluster 12 was significantly enriched in SSc but not OA and was highly enriched in immune terms related to antigen presentation and interferon-beta. Cluster 9 (*PLIN2+, MT1M+*) was significantly enriched in SSc, but not OA, and had a distinct metal ion signature with ontology terms associated with oxidative stress (“response to hypoxia”).

Taken together, our integrative analysis identified a large group of fibroblast subsets conserved between synovium and skin, with comparatively few tissue-specific clusters. Among conserved populations, disease-associated expansion involved progenitor cluster 1, immune-signaling clusters 3 and 6, fibrogenic clusters 5 and 8, and lining-like clusters 7 and 10, with distinct subsets enriched in OA, SSc, or both diseases. These findings suggest a strong overlap in disease-associated fibroblast activation states in OA synovium and SSc skin.

### Disease-enriched fibroblast subsets in OA and SSc converge on conserved fibrotic programs observed in idiopathic pulmonary fibrosis

To interrogate pro-fibrotic gene expression in OA synovium and SSc skin, we first defined a transcriptional “fibrotic template signature.” Idiopathic pulmonary fibrosis (IPF) is a high-mortality chronic disease defined by progressive lung fibrosis, leading to decline in lung function with premature death^32^. As a reference dataset for prototypic fibrotic disease, six scRNA-seq studies comprising healthy (n=54 subjects) and IPF lung (n=73 subjects) biopsies^33–38^ were identified and integrated (368,434 total cells, **Supp.-Fig.-4A-D**). Reactome PCA analysis identified fibroblasts as the most plastic cell type in fibrotic lung (**Supp.-Fig.-4E**), comparable to their high plasticity in OA synovium and SSc skin. Label transfer-based deconvolution confidently identified all fibroblast subsets from the integrated OA-SSc atlas in the lung except skin-specific cluster 11 (*COCH+, CRABP+*) (**Supp.-Fig.-4F, G**), suggesting strong parity in fibroblast states between OA synovium, SSc skin, and IPF lung. Myofibroblast-like cluster 5 (*SFRP2+, POSTN+*) and cluster 8 (*SFRP2+, COMP+*) were the most proportionally enriched subsets in IPF compared to healthy lung (*scProportionTest*, FDR < 0.05, |log2FD| > 0.58, **Supp.-Fig.-4H**), reinforcing the fibrogenic phenotype of these subsets.

We performed pseudobulk differential-expression analysis comparing IPF and healthy lung fibroblasts, yielding 247 upregulated and 244 downregulated robustly-differentially expressed genes (DEGs, *P_adj_* < 0.05, |log2FC| > 1, expression > 5% of fibroblasts in at least one condition), comprising a “pro-fibrotic gene signature.” Top upregulated DEGs in IPF fibroblasts included fibrosis- and ECM-associated genes such as *COL1A2*, *POSTN*, *COMP*, and *THY1* (**Fig.-3A**). Functional annotation confirmed a classical pro-fibrotic signature underpinning this gene set (**Fig.-3B**), including enrichment of “extracellular matrix organization”, “connective tissue development”, and “collagen fibril organization”. Intriguingly, osteochondral-related terms such as “ossification” and “cartilage development” were also enhanced.

**Figure 3:**
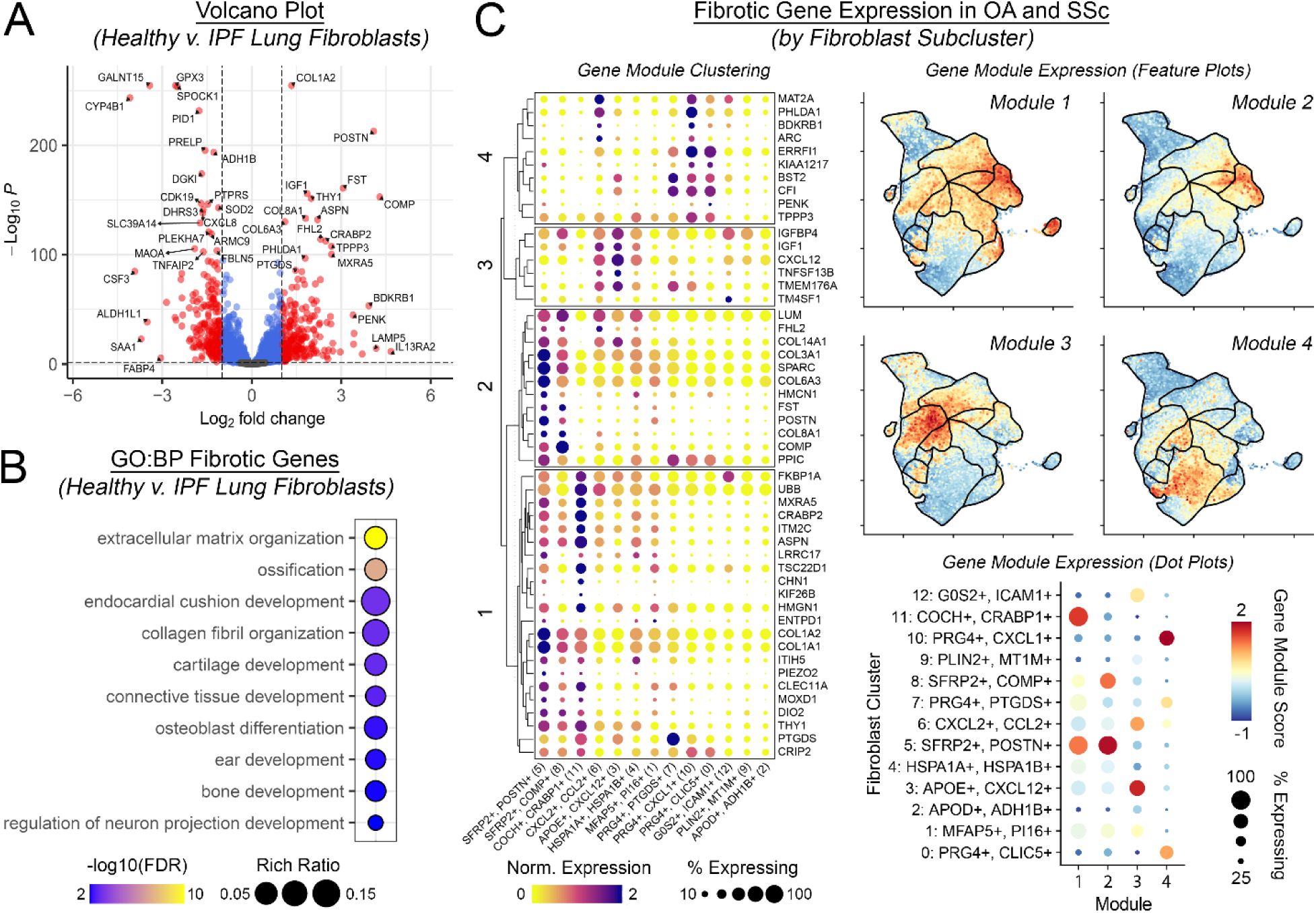
Disease-enriched fibroblast subsets in OA and SSc converge on conserved fibrotic programs observed in idiopathic pulmonary fibrosis. **(A)** Volcano plot of pseudobulk differential gene expression between IPF and healthy lung fibroblasts. Positive log2 fold-change values indicate increased expression in IPF compared to healthy lung. Differential expression was assessed using DESeq2 with a Wald test and *apeglm* log2 fold-change shrinkage; significant genes were defined by *P_adj_* < 0.05, |log2FC| > 1, and expression in >5% of fibroblasts in at least one condition. **(B)** Gene Ontology-Biological Process (GO:BP) enrichment analysis of genes significantly upregulated in IPF compared to healthy lung fibroblasts. Dot size indicates the rich ratio, and dot color indicates enrichment significance via PANTHER overrepresentation testing. **(C)** Mapping of the IPF-derived fibrotic gene signature onto the integrated synovium and skin fibroblast atlas. Left: Dot plot showing expression of the 50 most significantly upregulated IPF-derived genes across fibroblast subsets, organized into four gene modules by hierarchical clustering. Dot color represents normalized average expression, and dot size indicates the percentage of cells expressing each gene. Upper right: UMAP feature plots showing the distribution of module scores across integrated synovium and skin fibroblasts. Lower right: dot plot summarizing module expression across fibroblast subsets. Dot color represents the average module score, and dot size indicates the percentage of cells expressing genes within the module.

We mapped this fibrotic template gene signature onto synovium and skin fibroblasts via hierarchical clustering of the top 50 upregulated lung-derived fibrosis signature DEGs (sorted by –log10(*P_adj_*)). These genes were unbiasedly clustered into 4 distinct modules based on expression patterns in synovium/skin fibroblast subsets (**Fig.-3C**). Module 1 was expressed in myofibroblast-like cluster 5 (*SFRP2+, POSTN+*) and skin-specific cluster 11 (*COCH+, CRAPB+*) and was characterized by several markers of fibroblast activation and ECM production. Module 2 mapped more specifically to myofibroblast-like clusters 5 and 8 (*SFRP2+, COMP+*), comprised of ECM-associated genes (*LUM*, *POSTN*, *COMP*, and several collagens), which have been directly associated with disease-emergent myofibroblasts in both tissues^11,17^. Module 3 was expressed in immune-signaling clusters 3 (*APOE+, CXCL12+*) and 6 (*CXCL2+, CCL2+*) and consisted primarily of insulin-like growth-factor genes and chemokines (e.g. *IGF1*, *CXCL12*), reinforcing the chronic inflammatory component of fibrotic remodeling. Interestingly, module 4 mapped to lining and lining-like clusters, particularly 10 (*PRG4+, CXCL1+*), but also 0 (*PRG4+, CLIC5+*) and 7 (*PRG4+, PTGDS+*). In contrast to the fibrotic and ECM-related modules 1 and 2, module 4 comprised genes associated with epithelial differentiation, cellular activation, and tissue injury responses, including *BDKRB1, BST2, CFI, ERRFI1, PHLDA1,* and *PENK*.

Together, these findings demonstrate that transcriptomic hallmarks of prototypic fibrotic disease are highly enriched in conserved disease-associated fibroblast subsets in OA synovium and SSc skin.

### Disease-associated fibroblast subsets exhibit osteochondral-associated transcriptional programs

Ectopic mineralization and chondroid formation are well-known features of OA synovium and are also clinically observed in SSc skin. In OA, osteophyte formation has been linked to synovial activation and correlates strongly to patient-reported pain^4^. In SSc, ectopic mineralized tissue presents as calcinosis cutis^5^. Given the osteochondral signature observed in conserved, fibrosis-associated fibroblast subsets (**Fig.-2E**), we next characterized the extent of osteochondral signaling present in these cells.

Pathway enrichment of cluster-associated genes implicated progenitor fibroblast cluster 1 (*MFAP+, PI16+)* and fibrogenic clusters 5 (*SFRP2+, POSTN+*) and 8 (*SFRP2+, COMP+*) as enriched for several osteochondral terms including “chondrocyte development”, “cartilage development”, and “bone development” (**Fig.-4A**). Genes derived from these terms exhibited robust expression (up to 80% expression) within clusters 1, 5, and 8, with low-to-absent expression in lining-like and immune-signaling clusters (**Fig.-4B, Supp.-Fig.-5**). Disease-associated gene activation was observed in both OA synovium and SSc skin, including collagens (e.g. *COL1A1*, *COL3A1*, *COL6A1*), other ECM constituents (e.g. *LUM*, *OGN*, *BGN*, *ECM1*, *COMP*, *FBN1*, *FBLN5*), mediators of TGF (*TGFBI*) and WNT signaling (e.g. *CTNNB1*/β-catenin, *SFRP2*, *SFRP4*), and chondrogenic transcription factors (e.g. *RUNX1*, *NFIB*) (**Fig.-4C**). This is consistent with an osteochondral phenotype given the high induction of genes such as *COMP, LUM, SFRP2, THBS3,* and *BGN*. Many of these genes are also recognized fibrosis markers, suggesting an inherent overlap between these two processes. Gene-level expression of canonical chondrocyte markers (*ACAN*, *SOX9*) was low but detectable in all clusters with increased expression in synovium, however *COL2A1* expression was largely absent, suggesting the absence of *bona fide* chondrogenesis/mature chondrocytes in our data (**Fig.-4D**).

**Figure 4:**
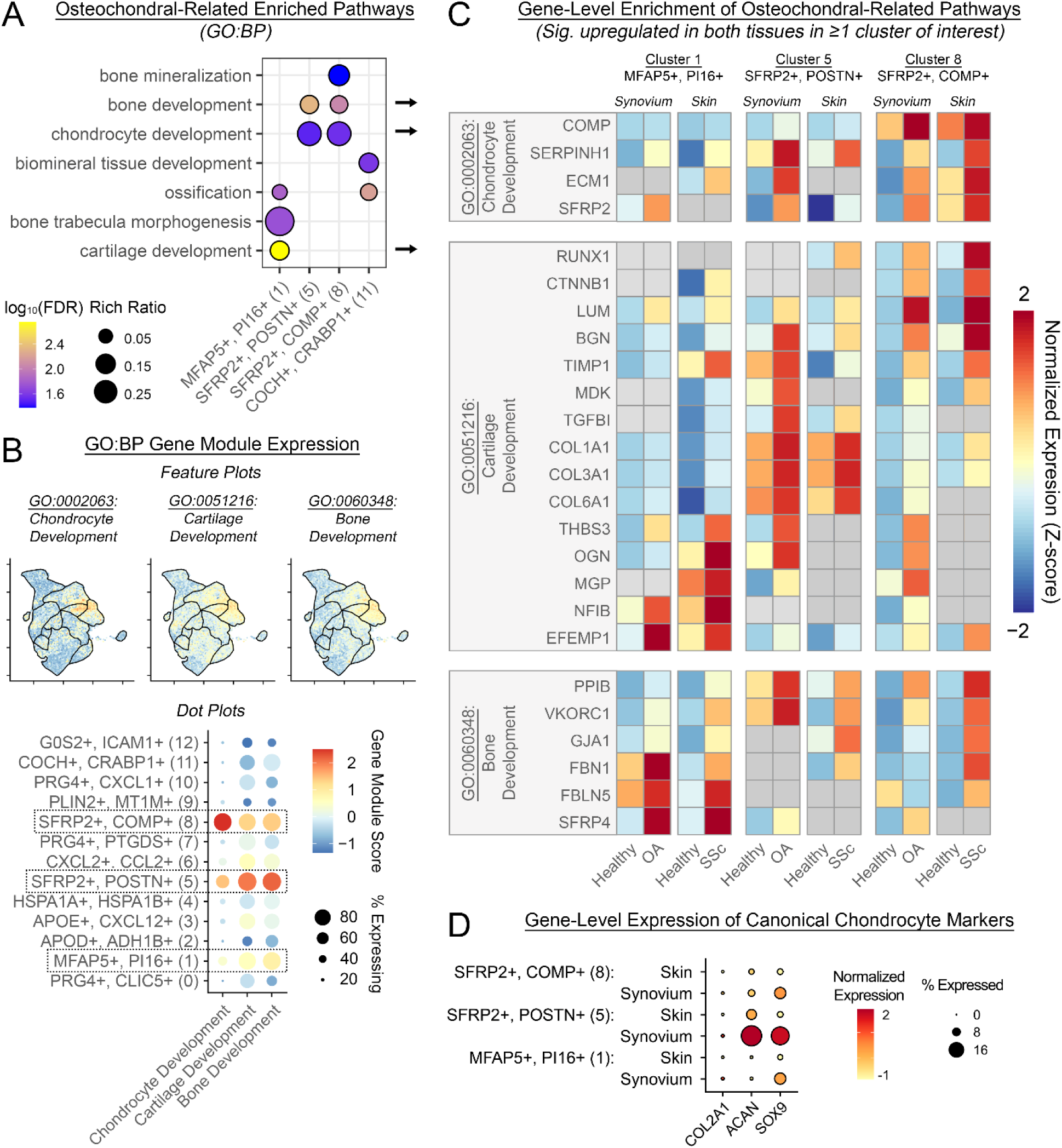
Disease-associated fibroblast subsets exhibit osteochondral-associated transcriptional programs. **(A)** Gene Ontology-Biological Process (GO:BP) enrichment analysis of cluster-defining genes showing selected osteochondral-related pathways enriched in fibroblast clusters 1 (*MFAP5+*, *PI16+*), 5 (*SFRP2+*, *POSTN+*), 8 (*SFRP2+*, *COMP+*), and 11 (*COCH+*, *CRABP1+*). Dot size indicates the rich ratio, and dot color indicates enrichment significance (−log10(FDR)) via PANTHER overrepresentation testing. **(B)** Expression of gene modules representing chondrocyte development (GO:0002063), cartilage development (GO:0051216), and bone development (GO:0060348) pathways in integrated synovium/skin fibroblast atlas. Top: UMAP feature plots of module scores. Bottom: Dot plot summarizing module expression across fibroblast subsets. Dot color indicates the average module score, and dot size indicates the percentage of cells expressing genes within each module. Boxes highlight clusters 1, 5, and 8, which exhibited coordinated enrichment of osteochondral programs. **(C)** Heatmaps showing gene-level expression of osteochondral pathway genes in clusters 1, 5, and 8, stratified by tissue and condition. Genes are grouped by GO:BP term, and synovium and skin expression values are scaled independently to illustrate healthy vs disease effects within each tissue. Color indicates normalized expression (Z-score); gray cells denote nonsignificant healthy vs disease comparisons. Only genes significantly upregulated in disease in both tissues and in at least one cluster of interest are shown. Genes represented in multiple pathways are displayed only under the first pathway in which they occur. **(D)** Dot plot showing expression of the canonical chondrocyte markers *COL2A1*, *ACAN*, and *SOX9* in clusters 1, 5, and 8, stratified by tissue. Dot color indicates normalized expression, and dot size indicates the percentage of cells expressing each gene.

Taken together, these findings show an active osteochondral gene program in fibrogenic disease-associated fibroblast subsets, suggesting transcriptional overlap between fibrotic signaling and osteochondral tissue formation.

### High-resolution spatial transcriptomics uncovers conserved spatial niches of disease-associated fibroblast subsets in OA and SSc

We next sought to define the spatial organization of disease-associated fibroblast subsets in OA synovium and SSc skin. Our analysis utilized Xenium 5K spatial transcriptomics, a subcellular-resolution, probe-based assay against 5000 mRNAs conducive to single-cell analysis. For SSc skin, we extracted three patients from a recent public dataset^15^. For OA synovium, we utilized archival FFPE blocks of human synovial biopsies (*n*=4) co-embedded into a tissue microarray. Cell types in each tissue dataset were clustered and annotated independently, and all expected major cell types were identified (**Supp.-Fig.-6A-B, 7A-B**). Fibroblasts underwent robust cell type deconvolution in each tissue independently, using the integrated OA-SSc fibroblast subsets as the reference, where >80% of OA Xenium fibroblasts and >65% of SSc Xenium fibroblasts were confidently mapped to one of the scRNA-seq fibroblast clusters (**Supp.-Fig.-6G, 7G**). Several fibroblast subsets (cluster 6, 7, 9-12) were not identifiable via deconvolution or had very low counts (**Supp.-Fig.-6C, 7C**), likely due to poor representation of their scRNA-seq-based gene profile in the Xenium probe set, combined with Xenium 5K’s lower detection depth insufficiently capturing low-expression genes (**Supp.-Fig.-6D-F, 7D-F**).

We focused primarily on myofibroblast-like clusters 5 (*SFRP2+, POSTN+*) and 8 (*SFRP2+, COMP+*), and immune-signaling cluster 3 (*APOE+, CXCL12+*), as pro-fibrotic clusters conserved across tissues which were consistently identifiable via Xenium. Spatial deconvolution identified two distinct anatomic niches which harbor these subsets in synovium and skin: 1) a fibrotic niche characterized by dense connective tissue, scarring, and fibrotic remodeling; and 2) a perivascular inflammatory niche, adjacent to endothelial cells, pericytes, and harboring pockets of immune infiltrate (**Fig.-5A-B**). In OA synovium, the fibrotic niche was represented in matrix-dense areas of the sub-synovium (**Fig.-5A**). The fibrotic niche in SSc skin was identified in areas of dense connective tissue and marked scarring (**Fig.-5B**). ECM-dense regions in SSc skin contained sparsely distributed fibroblasts from myofibroblast-like cluster 5 throughout the dermis, as compared to the densely packed OA synovium (**Supp.-Fig.-8A**). Cluster 8 in SSc skin was mostly localized to deeper layers of the dermis (**Supp.-Fig.-8B**), whereas it was more rarely observed in OA synovium. In both conditions, the fibrotic niche was populated more abundantly by cluster 5 than cluster 8, and their relative abundance (**Fig.-5C**) was comparable to that observed in scRNA-seq data (**Fig.-2C**).

**Figure 5:**
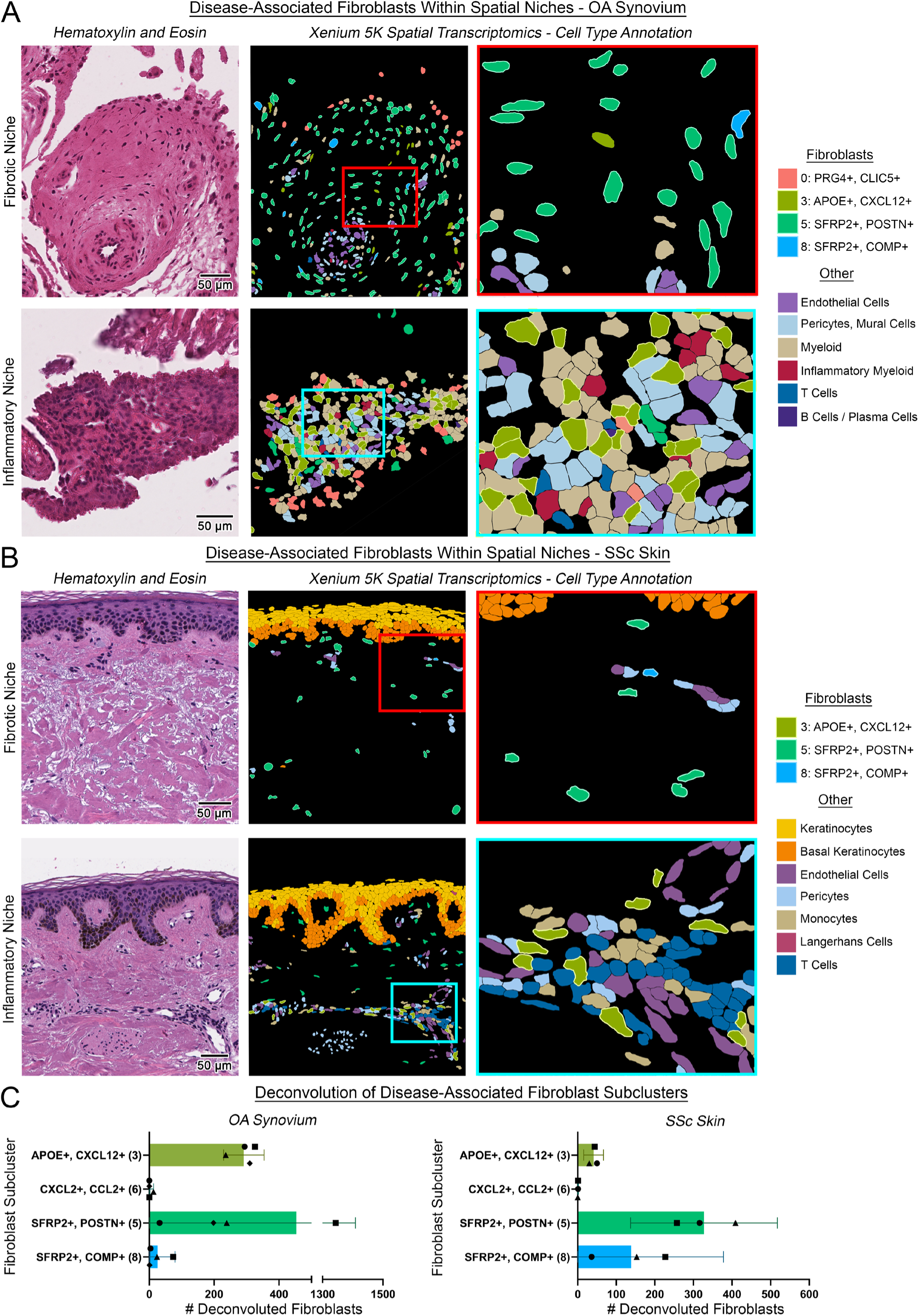
High-resolution spatial transcriptomics uncovers conserved spatial niches of disease-associated fibroblast subsets in OA and SSc. **(A-B)** Representative fibrotic and inflammatory niches in **(A)** OA synovium (fibrotic: OA_1; inflammatory: OA_3) and **(B)** SSc skin (fibrotic: SSc_6; inflammatory: SSc_2) via Xenium 5K spatial transcriptomics. Left: Hematoxylin and eosin-stained tissue sections. Center: Corresponding spatial transcriptomic maps showing cell-type annotations. Boxes indicate regions enlarged at right. Right: Enlarged views showing selected deconvoluted fibroblast subsets and neighboring stromal and immune cell types. Only fibroblasts assigned to a subcluster with a deconvolution probability ≥ 0.70 are shown. Scale bars = 50 *μm*. **(C)** Numbers of fibroblasts deconvoluted as disease-relevant clusters 3 (*APOE+, CXCL12+*), 5 (*SFRP2+*, *POSTN+*), 6 (*CXCL2+*, *CCL2+*), and 8 (*SFRP2+*, *COMP+*) in OA synovium and SSc skin Xenium samples. Bars indicate the mean, and error bars indicate 95% confidence intervals. Overlaid symbols represent individual subject samples.

The inflammatory niche harbored various immune cells surrounding blood vessels and capillaries identified morphologically by H&E, and endothelial cells and pericytes surrounded primarily by immune cells and fibroblasts identified by Xenium (**Fig.-5A-B**). The composition of immune infiltrates in the inflammatory niche differed between synovium and skin, with myeloid cells more abundant in synovium, while lymphocytes, namely T cells, were more abundant in the skin (**Supp.-Fig.-9**). This is concordant with cellular proportions determined by scRNA-seq (**Fig.-1C-F**) and potentially consistent with the prominent autoimmunity characteristic of SSc. Immune-signaling cluster 3 (*APOE+, CXCL12+*) was the primary subset identified within this niche (**Fig.-5A-B**), positioned at the interface between vasculature and various infiltrating immune cells, corroborating their proposed role in immunomodulation. Notably, in synovium, this niche was most prevalent around smaller capillaries close to the synovial lining, while deeper and larger vessels in the sub-synovium qualitatively exhibited lower abundance of cluster 3 fibroblasts (**Supp.-Fig.-9A**). This observation is consistent with prior work demonstrating the accumulation of immune cells through small, fenestrated capillaries below the synovial lining^39^.

Taken together, high-resolution spatial transcriptomics demonstrated that shared disease-associated fibroblast subsets occupy conserved microanatomical niches across OA synovium and SSc skin.

### Conserved fibroblast subsets exhibit overlapping disease-associated functional and regulatory programs

Having observed extensive similarities in transcriptional states and spatial organization of conserved disease-associated fibroblasts, we sought to assess whether these fibroblasts also exhibited shared disease-associated induction of biological pathways and underlying molecular regulators. We compared gene expression between healthy and diseased samples, focusing on the previously identified pro-fibrotic fibroblast subsets (**Fig.-3**) including myofibroblast-like clusters 5 (*SFRP2+, POSTN+*) and 8 (*SFRP2+, COMP+*), immune-signaling clusters 3 (*APOE+, CXCL12+*) and 6 (*CXCL2+, CCL2+*), and lining-like clusters 7 (*PRG4+, PTGDS+*) and cluster 10 (*PRG4+, CXCL1*+). Pseudobulk differential expression analysis defined significant DEGs between healthy and diseased fibroblasts stratified by tissue and cluster. Expectedly, both tissues exhibited unique sets of disease-associated gene expression - OA synovial fibroblasts exhibited this to a greater degree than SSc skin fibroblasts across clusters (**Fig.-6A, Supp.-Fig.-10A**), consistent with the greater degree of differential abundance observed between conditions in the synovium (**Fig.-2C**). Despite this, we measured marked overlap in disease-induced DEGs across fibroblast subsets of interest, with 33-55% of SSc DEGs also identified in OA (**Supp.-Fig.-10B**). The largest number of disease-induced DEGs (and highest DEG overlap in OA and SSc) was observed in myofibroblast-like clusters 5 (41% of SSc DEGs shared) and 8 (48% shared).

**Figure 6:**
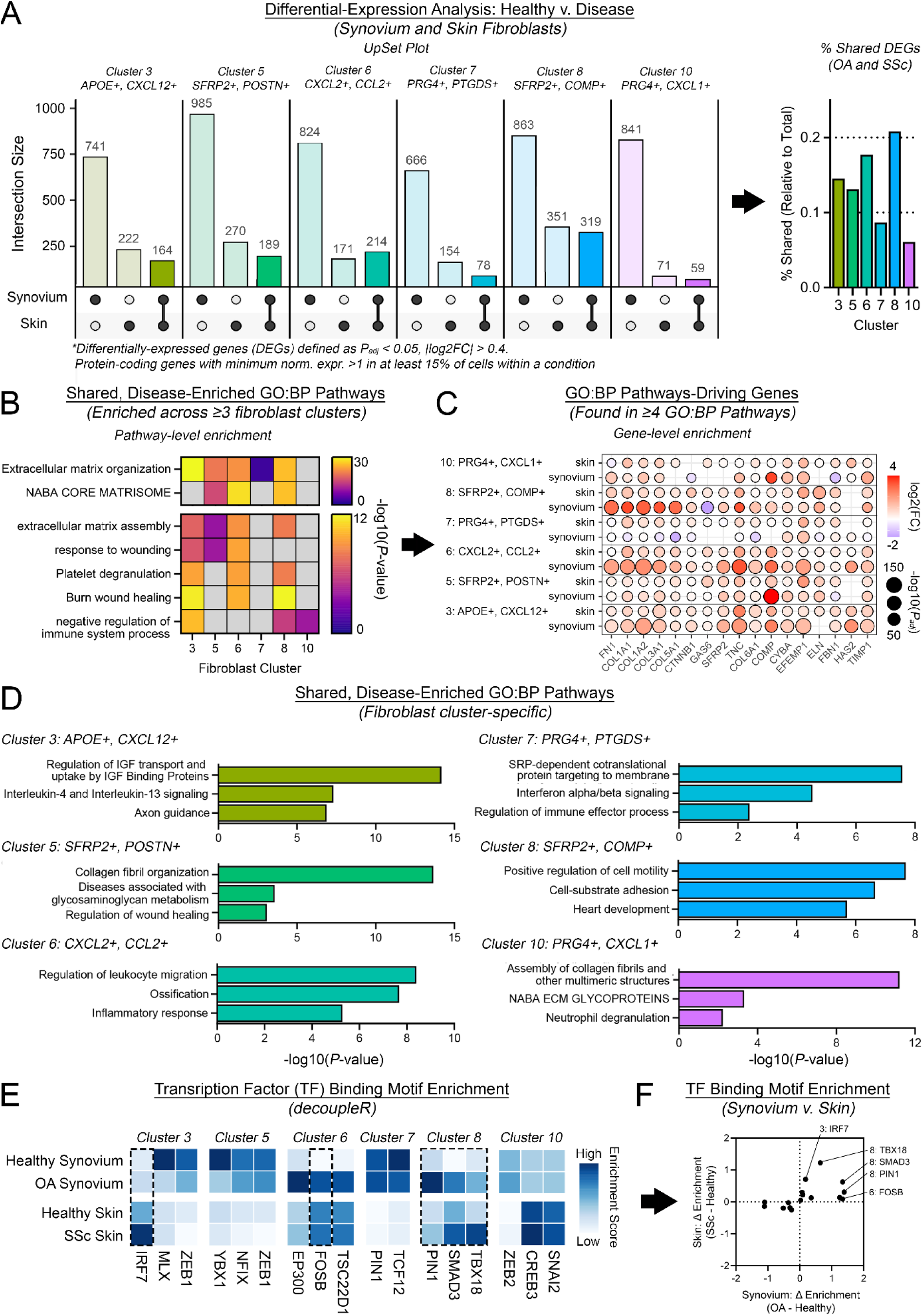
Conserved fibroblast subsets exhibit overlapping disease-associated functional and regulatory programs. **(A)** UpSet plots summarizing pseudobulk differential-expression analyses between diseased and healthy fibroblasts within synovium and skin for disease-associated clusters 3, 5, 6, 7, 8, and 10. Bars indicate the numbers of differentially expressed genes (DEGs) unique to OA synovium, unique to SSc skin, or shared between both diseases. DEGs were identified using DESeq2 with a Wald test and *apeglm* log2 fold-change shrinkage (*P*_adj_ < 0.05, |log2FC| > 0.4, and expression in >15% of cells in at least one condition). The bar graph on the right shows shared DEGs as a percentage of all SSc DEGs within each cluster. **(B)** Heatmap of shared disease-enriched Gene Ontology-Biological Process (GO:BP) pathways identified in at least three fibroblast clusters. Color indicates pathway-enrichment significance (−log10(*P*-value)), and gray denotes no significant enrichment. **(C)** Gene-level expression of genes contributing to at least four of the recurrent GO:BP pathways shown in (B), stratified by fibroblast cluster and tissue. Dot color indicates disease-associated log2FC, and dot size indicates significance (−log10(*P_adj_*)). **(D)** Selected shared disease-enriched GO:BP pathways specific to individual fibroblast clusters. Bars indicate enrichment significance (−log10(*P*-value)). **(E)** Heatmap of transcription-factor (TF) binding-motif enrichment inferred using *decoupleR* for healthy and diseased synovium and skin fibroblasts. For each cluster, the three TFs with the greatest average disease-associated change across synovium and skin are shown, restricted to TFs changing in the same direction in both tissues and expressed in >10% of cells in at least one condition in each tissue. Color indicates inferred TF enrichment score. **(F)** Comparison of disease-associated changes in inferred TF activity between OA synovium and SSc skin. Each point represents a cluster-specific TF. Positive values indicate increased activity in disease relative to healthy tissue. Selected TFs with concordant activation in both diseases are labeled.

Pathway analysis was used to functionally annotate shared DEGs in each cluster. Several pathways relating to ECM remodeling were enriched across 3 or more clusters (e.g. extracellular matrix organization and assembly) and wound healing (e.g. “response to wounding”, “burn wound healing”), suggesting that broad increases in these pro-fibrotic pathways occur across fibroblast clusters (**Fig.-6B**). Pathway-level enrichment was underpinned by conserved upregulation of genes related to ECM formation and remodeling (*FN1, COL1A1, COL3A1*), wound healing (*FBN1, SFRP2, COMP*), and immune-signaling pathways (*GAS6, CYBA*) (**Fig.-6C**). We also observed cluster-specific disease-associated enrichment of functional pathways (**Fig.-6D**). Clusters 3 (*APOE+, CXCL12+*) and 6 (*CXCL2+, CCL2+*) exhibited terms related to cytokine and chemokine signaling, with cluster 3 also implicating IGF signaling and neuronal remodeling (“axon guidance”), while “ossification” was enriched in cluster 6. In myofibroblast-like clusters, cluster 5 (*SFRP2+, POSTN+*) was enriched for additional terms related to matrix deposition and wound healing, reinforcing its myofibroblast-like identity, and cluster 8 (*SFRP2+, COMP+*) was enriched for terms such as “cell-substrate adhesion” and “positive regulation of cell motility”, consistent with the contractile nature of myofibroblast-like cells. Interestingly, both lining-like clusters were enriched for terms related to innate immunity (cluster 7 (*PRG4+, PTGDS+*): “Interferon alpha/beta signaling”, cluster 10 (*PRG4+, CXCL1*+): “neutrophil degranulation”), potentially implying an immune barrier role for these lining-associated fibroblasts.

Lastly, we asked whether these shared, disease-linked programs were associated with common upstream transcriptional regulators across OA and SSc. Activity scores were calculated to identify 3 TFs in each cluster with the highest average change between disease and healthy conditions in synovium and skin, prioritizing TFs which were differentially-activated in the same direction in both diseases (**Fig.-6E**). TFs without measurable expression in our data (>10% of cells in ≥1 condition in both tissues, **Supp.-Fig.10C**) were excluded. This analysis identified several TFs with consistently-enriched activity in both OA synovium and SSc skin. Of these, the most highly co-upregulated included IRF7 and FOSB in immune-signaling clusters 3 (*APOE+, CXCL12+*) and 6 (*CXCL2+, CCL2+*), respectively, and SMAD3, PIN1, and TBX18 in myofibroblast-like cluster 8 (*SFRP2+, COMP+*) (**Fig.-6F**).

Each of these transcription factors has been previously implicated in inflammatory or fibrotic signaling. IFR7 (Interferon Regulatory Factor 7) is a master regulator of inflammatory signaling, largely through type I interferon signaling. IFR7 has been implicated in driving IFN signaling in dermal fibroblasts in SSc^40^, macrophage polarization in rheumatoid arthritis (RA)^41^, and TNFα release in *in vitro* synovial fibroblast culture^42^. IFR7 has been shown to directly modulate several OA- and SSc-relevant inflammatory pathways including NFκB, cytokine/chemokine pathways interleukin and CXCL-, and prostaglandin signaling. FOSB (FosB proto-oncogene) is a component of the AP-1 transcription factor complex, along with the JUN, ATF, and MAF families, regulating gene transcription in response to a broad array of stimuli. In the context of disease-relevant inflammation and immune activation, FOSB+ fibroblasts have been identified in both RA synovium and SSc skin via spatial transcriptomics and are shown to be highly IFN and TNF-responsive^7,30^. Furthermore, FOSB has also been associated with pro-fibrotic macrophages^43^, suggesting this transcription factor may promote disease-associated inflammation across multiple cell types within the inflammatory niche.

Transcription factors SMAD3, PIN1, and TBX18 were enriched in OA and SSc in myofibroblast-like cluster 8 (*SFRP2+, COMP+*) and have been independently implicated in facilitating TGFβ signaling in the context of fibrosis. SMADs (including SMAD3) are a well-recognized family of signaling proteins crucial for TGFβ-driven ECM production and fibrotic activation, and SMAD3/TGFβ signaling in particular has been strongly linked to fibrosis^44^. PIN1 (Peptidyl-prolyl cis-trans isomerase NIMA-interacting 1) modulates phosphorylation, nuclear translocation, and transcriptional activation across multiple signaling pathways, and has been specifically shown to facilitate SMAD3/TGFβ signaling, SMAD/TGFβ signaling more broadly, and WNT/β-catenin signaling in the context of fibrosis^45–48^; thus, SMAD3 and PIN1 form a coordinated network of transcriptional control over SMAD/TGFβ signaling. TBX18 (T-box transcription factor 18) canonically regulates cardiovascular development and homeostasis^49^ and has been linked to cardiac and renal fibrosis, promoting fibrosis via TGFβ signaling^50,51^, though it has not been implicated in OA or SSc specifically.

Collectively, these findings demonstrate that fibroblast subsets conserved between OA synovium and SSc skin exhibit similar disease-associated transcriptional remodeling, characterized by convergent activation of ECM, wound healing, inflammatory, and immune-signaling pathways. These conserved gene programs are accompanied by overlapping predicted upstream transcriptional regulators, suggesting that common molecular mechanisms may underpin fibroblast activation across distinct fibrotic diseases and anatomic sites.

## Discussion

Fibroblasts are a highly plastic stromal cell capable of orchestrating fibrotic disease progression including ECM production, immunomodulation, and pro-angiogenic signaling. Here we employed a multi-omics approach to demonstrate the disease-associated shift in transcriptional phenotypes of human fibroblast subsets in OA and SSc, two diseases with divergent etiologies but multiple shared pathologic features. Our analyses revealed three major points of convergence between these two diseases: 1) Disease-associated activation of conserved fibroblast populations; 2) Spatial organization of these populations in analogous fibrotic and inflammatory tissue niches, and 3) Shared transcriptional programs putatively regulated by common upstream molecular mechanisms.

Cross-tissue single-cell atlases have established that multiple fibroblast subsets are conserved across tissues and diseases states^10,16,20^. Our analysis identified analogous progenitor, myofibroblast-like, immune-signaling, and lining-associated fibroblast populations in both OA synovium and SSc skin. This study extends prior work in three ways: it incorporates cells from both diseased and healthy tissues, demonstrates that conserved fibroblast subsets share disease-induced programs, and uses spatial transcriptomics to identify convergent anatomic niches containing the same disease-enriched fibroblast subsets in OA synovium and SSc skin. These findings further illustrate conserved disease mechanisms underlying these superficially divergent conditions.

Although excessive collagen deposition is the hallmark feature of fibrosis, chronic inflammation is also a well-established component of fibrotic disease^9,16,28,52^, and fibroblasts are increasingly recognized as regulators of inflammation by orchestrating the recruitment and activation of immune cells^9^. Inflammatory fibroblast populations expressing cytokines and chemokines have been identified by scRNA-seq analyses across multiple tissues and diseases, including the skin and synovium^10,11,16,52^, consistent with the transcriptional phenotypes of clusters 3 (*APOE+, CXCL12+*) and 6 (*CXCL2+, CCL2+*) identified within the inflammatory niche in our data. These clusters exhibited strong overlap in disease-enriched gene expression between OA and SSc, however, differences in the inflammatory niche were also apparent. Clusters 3 and 6 are significantly expanded in OA synovium, but present at comparable rates in healthy and SSc skin. Myeloid cells constitute the primary immune cell type within the synovium based on scRNA-seq and Xenium, while T cells were more abundant in skin. Despite these differences, immune-signaling fibroblasts in OA and SSc appear to promote a convergent phenotype of chronic inflammation and fibrosis, and we identified multiple transcription factors including IRF7 (cluster 3) and FOSB (cluster 6) enriched in both diseases. These are known mediators of multiple inflammatory pathways with established roles in both diseases, including IFN, TNF, multiple cytokine/chemokine pathways, and prostaglandin signaling. Further investigation is merited assessing their potential as target for immunomodulation via targeting these immune-modulating fibroblast populations.

Myofibroblasts are a well-characterized contractile and mechanosensitive fibroblast phenotype with an established, central role in fibrosis, persisting long-term in fibrotic tissue across multiple organ systems, and acting as the primary drivers of matrix formation and remodeling in injured tissue and during wound healing^8,17^. Xenium spatial transcriptomics identified these myofibroblast-like cells as the preeminent cells within a fibrotic niche characterized by high ECM deposition and scar tissue formation (within skin), reinforcing their role as direct drivers of ECM deposition, and substantial contributors to pro-fibrotic and osteochondral signaling-associated gene programs.

Furthermore, our data suggest that changes in enrichment and disease-induced changes in gene expression of myofibroblast-like clusters 5 (*SFRP2*+, *POSTN*+) and 8 (*SFRP2*+, *COMP*+) were highly conserved across OA and SSc fibroblasts, suggesting these cells represent a targetable opportunity for development of multi-disease fibrotic therapies. TF binding motif analysis identified 3 putatively shared transcription factors robustly activated in both diseases, with known roles in regulating TGFβ signaling. In particular, PIN1 and SMAD3 form a putative regulatory interaction driving conserved fibrogenic gene activation in myofibroblast-like cells in OA and SSc via TGFβ/SMAD signaling. PIN1 has been shown to impact the regulation of other key cellular processes and pathways as well^53^, including broader TGFβ/SMAD signaling and WNT/β-catenin^45–48^, in addition to SMAD3. Thus, PIN1 may represent an interesting target for further study due to its ability to modulate fibrogenic signaling across multiple disease-relevant signaling axes.

Spatial transcriptomics enabled us to confirm the *in situ* presence and contextualize the functional roles of disease-associated fibroblast subtypes predicted by scRNA-seq. However, this analysis is innately hindered by limitations of the technology. Certain fibroblast subclusters identified in the scRNA-seq data were challenging to detect via deconvolution in the spatial data, particularly as many top scRNA-seq based gene markers are absent from the Xenium 5K panel. This was especially challenging in rare or subtly different fibroblast subclusters such as immune-interacting clusters 3 (*APOE+, CXCL12+*) and 6 (*CXCL2+, CCL2+*), as well as lining-like clusters 0 (*PRG4+, CLIC5+*), 7 (*PRG4+, PTGDS+*), and 10 (*PRG4+, CXCL1*+). Further, the limited 2D nature of spatial transcriptomics may miss cells associated with rarer anatomical features.

The retrospective, meta-analytical nature of this data also introduces inherent limitations. Synovial and skin datasets were collected independently, without any possibility for matching patients by demographic factors, and biopsy timing differs between synovium and skin, with healthy skin biopsies being more commonly collected than healthy synovium. Within-tissue differences in biopsy location, processing pipeline, inclusion criteria, and sample number introduced further variability between studies. We attempted to mitigate these limitations and achieve robust alignment in scRNA-seq data via rigorous data integration, scaling, and normalization^54^. Cell counts also varied on the basis of tissue, disease, study, and clustering, potentially driving sample size-dependent effects. As the overall incidence of the various fibroblast clusters within individual subjects is insufficient for cluster-level pseudobulk analysis utilizing individual subjects as controls, we employed the DElegate R package for pseudobulk differential-expression analyses, which creates pseudo-replicates within each condition, normalizing for input cell number^55,56^. Our fibrotic gene signature, derived from a comparison of IPF and healthy lung tissue, did not consider other fibrotic diseases such as kidney or liver fibrosis and is therefore inherently biased towards lung fibrosis, as the availability of matched healthy and diseased scRNA-seq data in these diseases was limited at the time of our query.

This study demonstrates that OA and SSc exhibit conserved fibroblast subsets, occupying analogous spatial niches and exhibiting convergent disease-associated activation programs governed by common putative molecular mechanisms. This suggests that fibroblast-mediated fibrosis may be governed by common stromal principles across anatomically- and etiologically-distinct diseases. By defining the cellular, spatial, and molecular features shared between OA synovium and SSc skin, this study contributes to development of multi-disease anti-fibrotic therapeutic strategies and advances a growing view of fibrosis as conserved pathological program, rather than a tissue-specific disease manifestation.

## Methods

### OA Synovium H&E Image

Synovial tissue was formalin-fixed, paraffin-embedded, sectioned at 5-μm thickness, and stained with hematoxylin and eosin (H&E) using standard histopathologic methods. The photomicrograph (H&E-stained, x6 magnification) of a representative region of synovial fibrosis was provided by an expert musculoskeletal pathologist (E.D.). The fibrosis appears as a bandlike zone of connective tissue beneath the synovial lining layer associated with loss of the regional adipose tissue.

### SSc Skin H&E Image

3mm punch biopsies of lesional (SSc) skin were obtained. Tissue biopsies were formalin-fixed and paraffin embedded. 3-5 um thick tissue sections were processed for routine histological staining with hematoxylin and eosin (H&E).

### Data Acquisition

Publicly available scRNA-seq datasets were downloaded from the Gene Expression Omnibus database (GEO) across human OA synovium, SSc skin, IPF and their healthy counterparts, when available. 5 synovium datasets^12,18,22–24^, 4 skin datasets^14,25–27^, and 6 lung datasets^33–38^ were identified.

### Quality Control

Quality control was performed computationally in R (v4.4.0) using RStudio (v2024.12.1) and *Seurat*^57^ (v5.1.0). For studies that provided raw feature barcode matrices, *DropletUtils*^58,59^ (v1.27.2) was used to produce filtered feature barcode matrices by identifying empty droplets in the raw data. Cells in each sample were then filtered to preserve only high quality, viable cells: cells with mitochondrial gene ratio > 10% were removed to exclude non-viable cells^60^, cells with greater than the 93^rd^ percentile of features (nFeature > 93%) were removed to exclude multiplets, and cells with less than 200 unique genes (nFeature < 200) or with less than 1800 UMI (nCount < 1800) for synovium or 800 UMI (nCount < 800) for skin and lung were excluded to remove debris and other low quality droplets - these cutoffs were determined empirically for each dataset by iteratively assessing a broad range of cutoffs to determine the optimal point which excluded low-quality droplets while preserving as many valid cells as possible.

### Integration and Clustering (All Cells)

Datasets for synovium and skin were each integrated by study using SCTransform normalization and scaling, followed by RPCA-based integration using S*eurat*. Linear and non-linear dimensionality reduction were performed using principal component analysis (PCA) and uniform manifold approximation and projection (UMAP), respectively. Dimensions and resolutions for UMAP and clustering were determined based on elbow plot results from principal component analysis and analysis of cluster-defining markers (Synovium: dimension=35, resolution=0.25; Skin: dimension=31, resolution=0.1). FindAllMarkers() was used for cell type annotation in each tissue atlas.

Sketch-based integration in *Seurat* was used for the lung dataset to accommodate the high cellularity of the data and offload computational resources. 5000 cells were randomly sampled from all lung studies to create the sketched dataset. Data was integrated by study, using log normalization and RPCA integration (k.anchors=30). Dimensionality reduction was performed on the sketched data (dimension=30, resolution=0.25). The full dataset was then projected onto the sketched data to produce the full UMAP. FindAllMarkers() was used for cell type annotation based on markers.

### Reactome PCA Analysis

The R package *ReactomeGSA*^61^ (v1.22.0) was used to perform gene set variance analysis and PCA of healthy and diseased cells in each cluster, using the Reactome pathways database. The analysis was run on each tissue atlas (synovium, skin, and lung) individually using default settings. Euclidean distances were calculated from the healthy cells in the cluster to its disease counterpart. Enrichment scores for pathways that were positively enriched in both OA synovium and SSc skin compared to healthy controls were then curated to generate a heatmap of disease-relevant co-enriched signaling pathways.

### Fibroblast Integration and Clustering

Fibroblasts were identified using positive expression of *PDGFRA* and negative expression of *PTPRC/CD45*. Fibroblasts were then subset from the synovium and skin atlases and integrated together using SCTransform normalization and RPCA integration. Clustering dimensions and resolutions were picked based on PCA and marker analysis (dimensions=25, resolution= 0.2). Unsupervised clustering initially produced 14 clusters (0–13). Marker gene analysis revealed one cluster comprised of fibroblast-endothelial and fibroblast-myeloid multiplets, co-expressing canonical fibroblast markers with myeloid markers *FOLR2*, *CD163*, *ITGAM*, and *LYVE1*, and endothelial markers such as *PECAM1* and *VWF,* which were not detected in the other fibroblast clusters. This cluster was thus determined to contain heterogeneous multiplets, and was removed from the dataset, and we proceeded with 13 clusters for downstream analysis.

### Fibroblast Subcluster Annotations

FindAllMarkers() was used to generate differentially expressed genes for each of the clusters. Markers were filtered based on their level of expression (expressed in > 50% of cells in the cluster, and less than 50% of all other cells), then sorted by log2fc to identify each cluster by 2 defining markers to refer to each cluster by. In cases where clusters shared key gene markers (e.g. *SFRP2* denoting myofibroblast-like cells and *PRG4* denoting synovial lining-like cells), these markers were used for all relevant clusters to denote this shared feature, with a second unique marker used to delineate between them. Final cluster defining genes were as follows: Cluster 0 (*PRG4+*, *CLIC5+*), Cluster 1 (*MFAP5+*, *PI16+*), Cluster 2 (*APOD+*, *ADH1B+*), Cluster 3 (*APOE+*, *CXCL12+*), Cluster 4 (*HSPA1A+*, *HSPA1B+*), Cluster 5 (*SFRP2+*, *POSTN+*), Cluster 6 (*CXCL2+*, *CCL2+*), Cluster 7 (*PRG4+*, *PTGDS+*), Cluster 8 (*SFRP2+*, *COMP+*), Cluster 9 (*PLIN2+*, *MT1M+*), Cluster 10 (*PRG4+*, *CXCL1+*), Cluster 11 (*COCH+*, *CRABP1+*), and Cluster 12 *(G0S2+*, *ICAM1+)*.

### Fibroblast Subcluster Analysis

Abundances of healthy and diseased cells (by tissue) were calculated, reporting both absolute counts and normalized fractions of total. *scProportionTest*^62^ (v0.0.0.9000) was used to statistically test for clusters which were significantly enriched in disease by using a permutation test to calculate p-values for each cluster and then uses bootstrapping to calculate a confidence interval for difference in magnitude between conditions (FDR < 0.05, |log2FD| > 0.58). Differentially expressed genes (DEGs) for each cluster were determined using FindAllMarkers() in Seurat (*P_adj_* < 0.05, positive log2fc, expressed in > 50% of cells in the cluster). Final filtered gene lists per cluster were submitted to PantherDB^63,64^ for statistical overrepresentation testing using the Gene Ontology Biological Processes (GO:BP) database. *decoupleR*^65^ (v2.14.0) was used to calculate mean pathway activity scores across all fibroblast subclusters using the PROGENy pathway database. A multivariate linear model was used to calculate the enrichment scores for each pathway, fitting a linear model to predict gene expression based on weights of the pathway-gene interactions for all pathways. T-values of the slopes of the linear model are reported.

### Chondrogenic Fibroblast Module

GO:BP terms related to cartilage and bone development were identified from previously submitted gene lists. Gene module scores were calculated using AddModuleScore() with genes that were upregulated in both synovium and skin and expressed in at least one of the three clusters of interest. Z-scores were calculated for genes comprising the modules for each term. Synovium and skin expression are scaled within tissue. Only significant disease comparisons are shown (*P_adj_* < 0.05).

### Lung Fibrosis Module

Pseudobulk differential expression analysis with using *DElegate*^55,56^ (v1.2.1) was performed using the DESeq2 algorithm (Wald test) with *apeglm* log2fc shrinkage, to determine differential gene expression between healthy and IPF fibroblasts (*P_adj_* < 0.05, |log2fc| > 1, rate of expression > 5% in at least one condition). The *apeglm* method adjusts effect sizes based on gene information, shrinking large effects of low-throughput genes, and preserving effects of high-throughput genes. Disease-enriched genes (log2fc > 1) were submitted to PantherDB^63,64^ for statistical overrepresentation test using the GO:BP database. The top 50 upregulated DEGs (sorted by –log10(*P_adj_*)) were isolated, and their expression within synovium/skin fibroblast subclusters was assessed to determine which clusters were the most pro-fibrotic. These 50 DEGs were clustered into gene modules using *scCustomize*^66^ (v2.1.2) based on their expression within synovium/skin fibroblast subclusters. Initially, no pre-defined number of clusters (*k*) was set, but based on review of the preliminary clustering tree and associated elbow plot, *k*=4 was selected for final gene module clustering as presented in main text Figure 3. To visualize gene module expression within synovium/skin fibroblast subclusters, gene module scores were calculated using AddModuleScore() and visualized via dot and feature plots in *Seurat*^57^. Cell type deconvolution of the lung fibroblasts was performed using the MapQuery() function in *Seurat*, using the integrated OA and SSc fibroblasts as a reference.

### Spatial Transcriptomics

Publicly available Xenium 5K Prime spatial transcriptomics data of SSc skin were downloaded from GEO (GSE312932)^15^. Three samples were selected per tissue (on the basis of high section quality), merged computationally using *Seurat* and clustered to resolve broad cell types (dim=35, res=0.4), which were annotated using markers. Low-quality cells clustered together (nCount < 250, no upregulated gene markers) and were removed from the dataset using the lasso tool.

Osteoarthritis synovium samples were collected from archival FFPE blocks at University Hospital Zurich (USZ). Tissues were obtained during joint replacement or ultrasound-guided synovial biopsy. Adjacent H&E-stained sections were used to determine ROI cores that were isolated from blocks and re-embedded into a single block. Samples were sectioned for spatial transcriptomics using 10x Genomics Xenium 5K platform (Human Pan Tissue & Pathways Panel with 50 custom gene panel) and processed at the Functional Genomics Center Zurich. Raw Xenium data was loaded in R (v4.4.3) using *Seurat*^57^ (v5.3.0). Samples were computationally isolated via ROI selection. Low-quality cells were removed per sample by thresholding on minimum transcript and gene count, an upper transcript-count bound at the 99th percentile, absence of control or blank codeword counts, and a nucleus-area window (4–80 µm²). Count/feature thresholds were tuned per sample (minimum 10–40 transcripts and 5–20 genes). QC-filtered Seurat objects were normalized per sample and merged. Expression data was scaled, regressing total transcript counts per cell. Principal component analysis was performed, and the first 33 PCs were used for downstream analysis. Batch effects across samples were corrected with Harmony (v1.2.4; ʎ=3). Louvain clustering was performed on the Harmony embedding and clusters were manually annotated into cell types based on marker genes. Low-quality clusters were excluded from downstream analyses.

In both Xenium datasets, the fibroblasts were identified and cell type deconvolution was performed using *InSituType*^67^ (v2.0), with the integrated OA-SSc fibroblast object as the reference. Only cells pertaining to the disease were used for the deconvolution for the corresponding Xenium dataset. Fibroblasts with a deconvolution posterior probability of ≥ 0.7 were assigned to that fibroblast subcluster annotation. Any fibroblast that did not pass this threshold retained the label of Fibroblast and is not shown in visualization and is excluded from quantifications. All other cell types retained their original annotation.

### Synovium and Skin Fibroblast Differential Expression Analysis

Pseudobulk differential expression analysis using *DElegate*^55,56^ was performed using the DESeq2 algorithm (Wald test) with *apeglm* log2fc shrinkage, to determine differential gene expression between fibroblasts from healthy and disease (OA/SSc) groups, within each tissue and fibroblast subcluster. Because cells from individual subjects generally did not possess sufficient cellularity within each fibroblast subcluster to enable subjects to be used as bio-replicates, fibroblasts within each condition were randomly assigned into pseudo-replicates (*n*=3 reps/condition) within each cluster, using *DElegate*. The resultant gene lists were filtered to include only significantly upregulated genes with meaningful expression within a cluster (*P_adj_* < 0.05, |log2fc| > 0.4, rate of expression > 15% in at least one condition within a cluster) and sorted into the following categories: unique to synovium, unique to skin, or overlapping between tissues. *UpSetR*^68^ (v1.4.0) was used to visualize the set sizes and their intersection for disease-enriched, pro-fibrotic clusters 3, 5, 6, 7, 8, and 10 as UpSet plots. The intersecting gene lists were filtered by positive log2fc to isolate positively enriched genes and submitted to Metascape^69^ for pathway analysis, for each cluster individually. Terms overlapping in ≥ 3 clusters were identified. Overlapping genes in these terms were identified, and gene level expression was shown.

Transcription factor (TF) binding motif enrichment was performed using *decoupleR*^65^, which employs the collectTRI database to calculate enrichment scores for each transcription factor. TF enrichment scores are inferred using a multivariate linear model, which fits a linear model based on the TF-gene interaction weights to predict gene expression. T-values of the slopes are reported. The top 50 predicted TFs were extracted for each cluster and filtered to select TFs that changed in the same direction in both tissues. These TFs were then filtered to genes expressed in > 10% of all cells in a cluster in the conditions with the greater enrichment score (either both healthy condition or both diseased conditions). The top 3 TFs, ranked by average magnitude of change, were visualized. The difference between the disease and healthy scores in both tissues were plotted in a scatter plot to determine the most differentially perturbed TFs shared across OA and SSc.

### Plotting and Visualization

Heatmaps were generated using *pheatmap*^70^ (v1.0.13). Volcano plots were generated using *EnhancedVolcano*^71^ (v1.26.0). Feature Plots, UMAPs, Dot Plots, Gene Modules, and Xenium ROIs were generated using *Seurat*^57^. Bubble plots were generated using *ggplot2*^72^ (v4.0.1). Clustered dot plots for fibrotic gene module analysis were generated using *scCustomize*^66^ (v2.1.2). All other plots were generated using GraphPad Prism (v10.3.0). Tables were generated in Microsoft Excel.

## Data Availability Statement

The following data were obtained from previously published studies cited in this article: Synovium scRNA-seq datasets can be found on the Gene Expression Omnibus (GEO) at the following accession numbers: GSE283080, GSE152805, GSE246416, GSE233502, and GSE126651. Skin scRNA-seq datasets can be found on GEO at the following accession numbers: GSE214088, GSE181957, GSE138669, and GSE160536. Lung scRNA-seq datasets can be found on GEO at the following accession numbers: GSE156310, GSE128033, GSE136831, GSE135893, GSE143706, and GSE122960. Systemic sclerosis skin Xenium dataset can be found on GEO at the accession GSE312932. Newly-generated human OA Xenium data is available upon reasonable request.

## Author Contribution Statement

A.M. and M.D.N. contributed to conceptualization, methodology, investigation, visualization and writing. N.B. and A.K. contributed to investigation, visualization, and writing. T.M.H. and C.O. contributed to investigation and writing. E.D. contributed to visualization. D.O., J.V., and T.M. contributed to conceptualization, methodology, and writing.

## Funding Statement

T.M. – National Institutes of Health / National Institute of Arthritis and Musculoskeletal and Skin Diseases (R01 AR080035, UC2 AR082186)

J.V. - NIH, DoD, Rheumatology research foundation

C.O. - EU Project MDR RA Grant Number: 101155807

D.O. - RE-JOIN consortium NIH-NIAMS UC2AR082186

