## Supplemental Figures for "Conserved Fibroblast Subsets Drive Shared Fibrotic Mechanisms in Osteoarthritis Synovium and Systemic Sclerosis Skin"

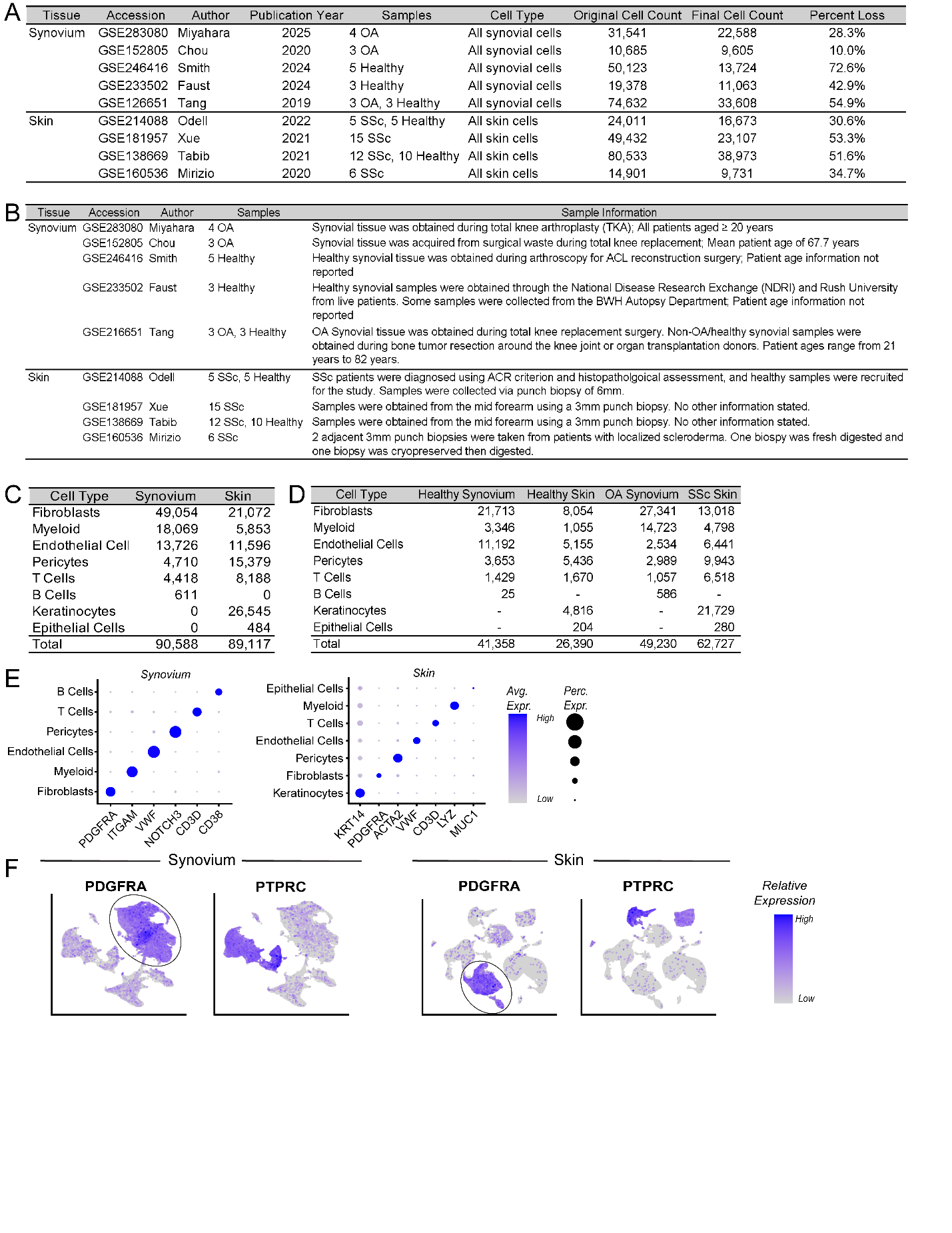

### Supplemental Figure 1: Study information and cell type identification

**(A)** Tabulated study summaries of synovium and skin scRNA-seq datasets included in the analysis. “Original Cell Count” represents the cell count in raw data, and “Final Cell Count” represents cells included in analyses following quality control. “Percent Loss” represents the proportion of cells removed during quality control. **(B)** Sample collection and available clinical information for each included dataset. **(C)** Cell counts by major cell type in synovium and skin scRNA-seq integrations. **(D)** Cell counts by major cell type and by disease condition in scRNA-seq integrations for synovium and skin. **(E)** Dot plots of canonical marker-gene expression used to annotate major cell types in the synovium and skin datasets. Dot color indicates average scaled expression, and dot size indicates the percentage of cells expressing each gene. **(F)** UMAP feature plots showing relative expression of *PDGFRA* and *PTPRC* in the integrated synovium and skin datasets. Circled regions indicate the *PDGFRA*+, *PTPRC*- populations identified as fibroblasts and retained for downstream analysis.

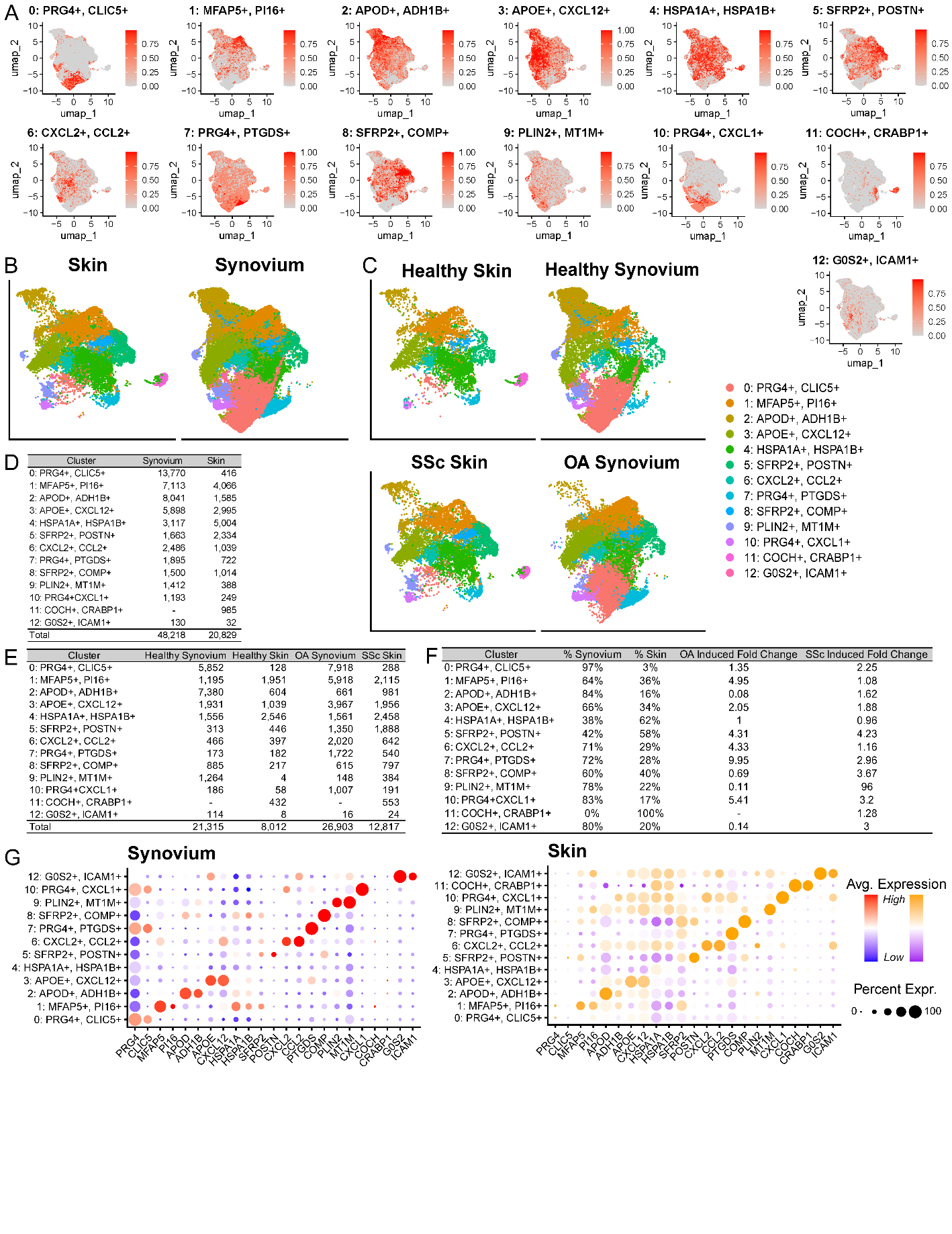

### Supplemental Figure 2: Fibroblast subset identification and proportions

**(A)** UMAP feature plots showing gene module scores of cluster-defining marker genes across fibroblast subsets identified in the synovium and skin integration **(B)** Integrated fibroblast UMAPs split by tissue. **(C)** Integrated fibroblast UMAPs split by tissue and disease condition. Colors in (B–C) indicate cluster identity based on annotation shown on right. **(D)** Fibroblast counts across clusters, stratified by tissue. **(E)** Fibroblast counts across clusters, stratified by tissue and disease condition. **(F)** Tissue distribution and disease-associated changes in fibroblast-cluster abundance. Percentages indicate the relative contribution of synovium and skin to each cluster after normalization for total tissue-specific fibroblast counts. Fold changes indicate cluster abundance in each disease condition relative to the respective healthy condition. **(G)** Dot plots showing expression of cluster-defining marker genes across fibroblast subsets in synovium (left) and skin (right). Dot color indicates scaled average expression within each tissue, and dot size indicates the percentage of cells expressing each gene.

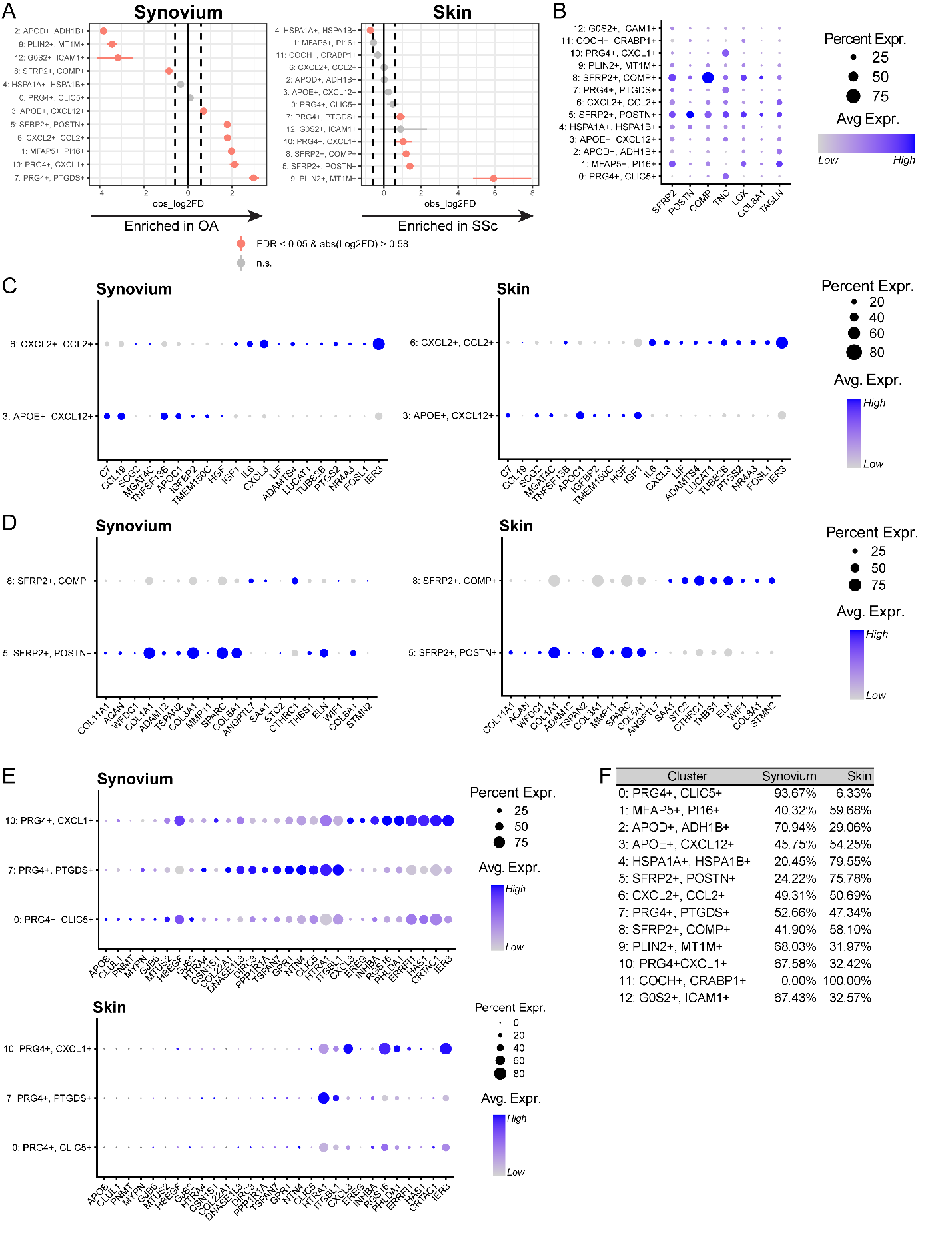

### Supplemental Figure 3: Further characterization of fibroblast subsets

**(A)** Forest plots of differences in cluster abundance across fibroblast subsets between diseased and healthy conditions in synovium and skin, by *scProportionTest*. Points indicate observed log2FD, and horizontal lines indicate bootstrapped confidence intervals. Red denotes significantly altered clusters (FDR < 0.05 and |log2FD| > 0.58) and gray denotes nonsignificant comparisons. **(B)** Dot plot showing expression of canonical myofibroblast-associated genes across fibroblast clusters. **(C–E)** Dot plots comparing the expression of cluster-defining genes among transcriptionally-related fibroblast subsets in synovium and skin: **(C)** immune-signaling clusters 3 (*APOE+, CXCL12+*) and 6 (*CXCL2+, CCL2+*); **(D)** myofibroblast-like clusters 5 (*SFRP2+, POSTN+*) and 8 (*SFRP2+, COMP+*); and **(E)** lining and lining-like clusters 0 (*PRG4+, CLIC5+*), 7 (*PRG4+, PTGDS+*), and 10 (*PRG4+, CXCL1+*). Expression profiles demonstrate distinct gene-expression signatures among related clusters and tissue-dependent expression of selected markers. In (B–E), dot color indicates scaled average expression, and dot size indicates the percentage of cells expressing each gene. **(F)** Relative contribution of synovium and skin to each fibroblast cluster after normalization for total tissue-specific fibroblast counts.

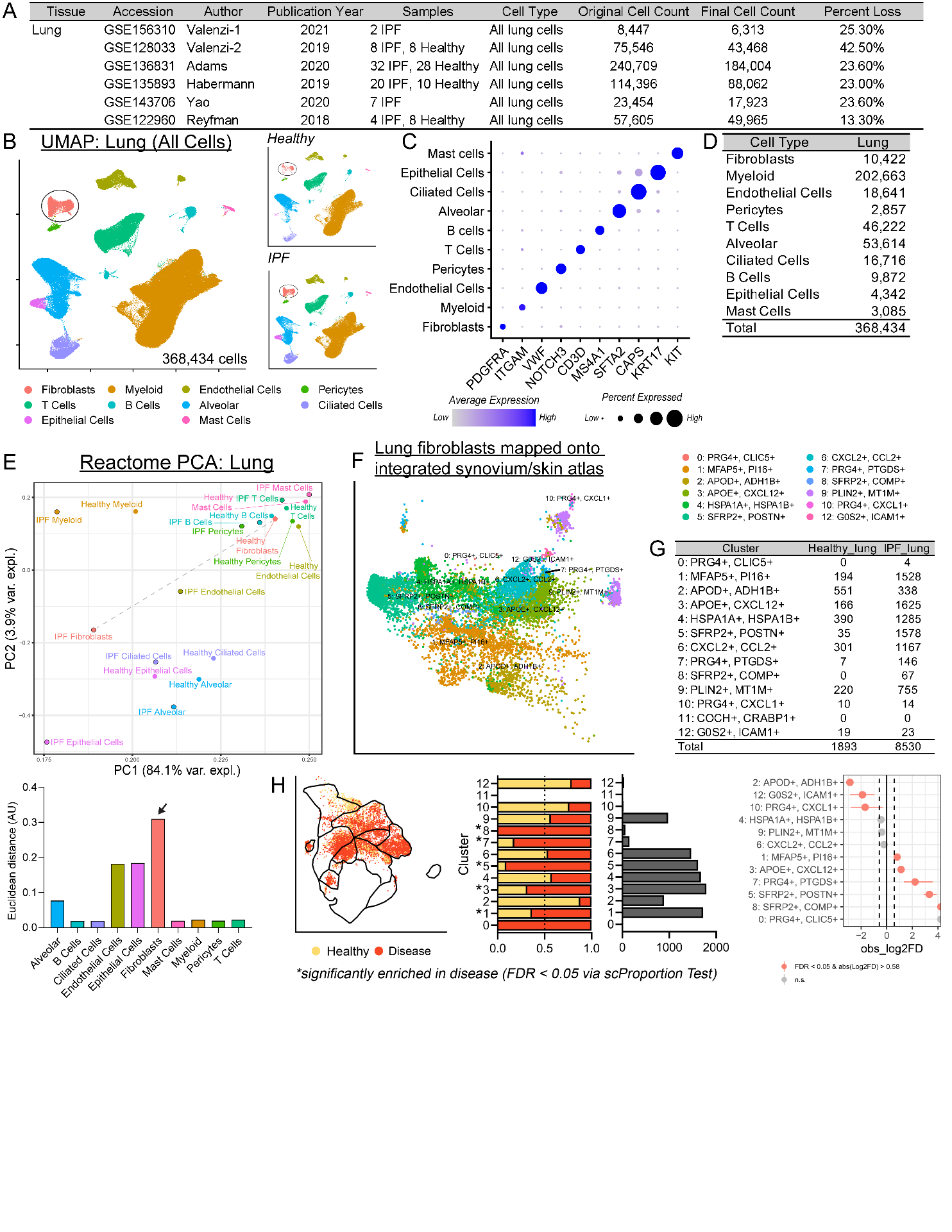

### Supplemental Figure 4: Integrated healthy and idiopathic pulmonary fibrosis lung atlas and cross-tissue mapping of fibroblast subsets.

**(A)** Tabulated study summaries of lung scRNA-seq datasets included in the analysis, and total cell counts before and after quality control. **(B)** UMAP of 368,434 cells integrated across healthy and idiopathic pulmonary fibrosis (IPF) lung datasets, annotated by major cell type. Fibroblasts retained for downstream analysis are circled. **(C)** Dot plot showing canonical marker-gene expression across major lung cell types. Dot color indicates average expression, and dot size indicates the percentage of cells expressing each gene. **(D)** Cell counts of major cell types in the integrated lung atlas. **(E)** Principal component analysis (PCA) of Reactome pathway enrichment profiles for major cell types in healthy and IPF lung. Top: Points on the PCA plot represent cell type and healthy/disease condition. Bottom: Euclidean distances between matched healthy and diseased cell types quantify the magnitude of disease-associated functional divergence. **(F)** UMAP of lung fibroblasts after label transfer of fibroblast-cluster annotations from the integrated synovium and skin fibroblast atlas. Colors indicate transferred fibroblast-cluster identity. **(G)** Fibroblast cell counts for each transferred cluster stratified by healthy and IPF lung. **(H)** Relative abundance of transferred fibroblast clusters in healthy and IPF lung. Left: Healthy and IPF lung fibroblasts mapped onto UMAP space of integrated synovium/skin fibroblast atlas. Center: Stacked bars show the normalized fraction of healthy and IPF cells within each cluster, and adjacent bars show absolute cell counts. Right: Forest plot showing differences in cluster abundance between IPF and healthy lung, as determined by *scProportionTest*. Points indicate observed log2FD, and horizontal lines indicate bootstrapped confidence intervals. Red denotes significantly altered clusters (FDR < 0.05 and |log2FD| > 0.58), gray denotes nonsignificant comparisons, and asterisks indicate clusters significantly enriched in IPF.

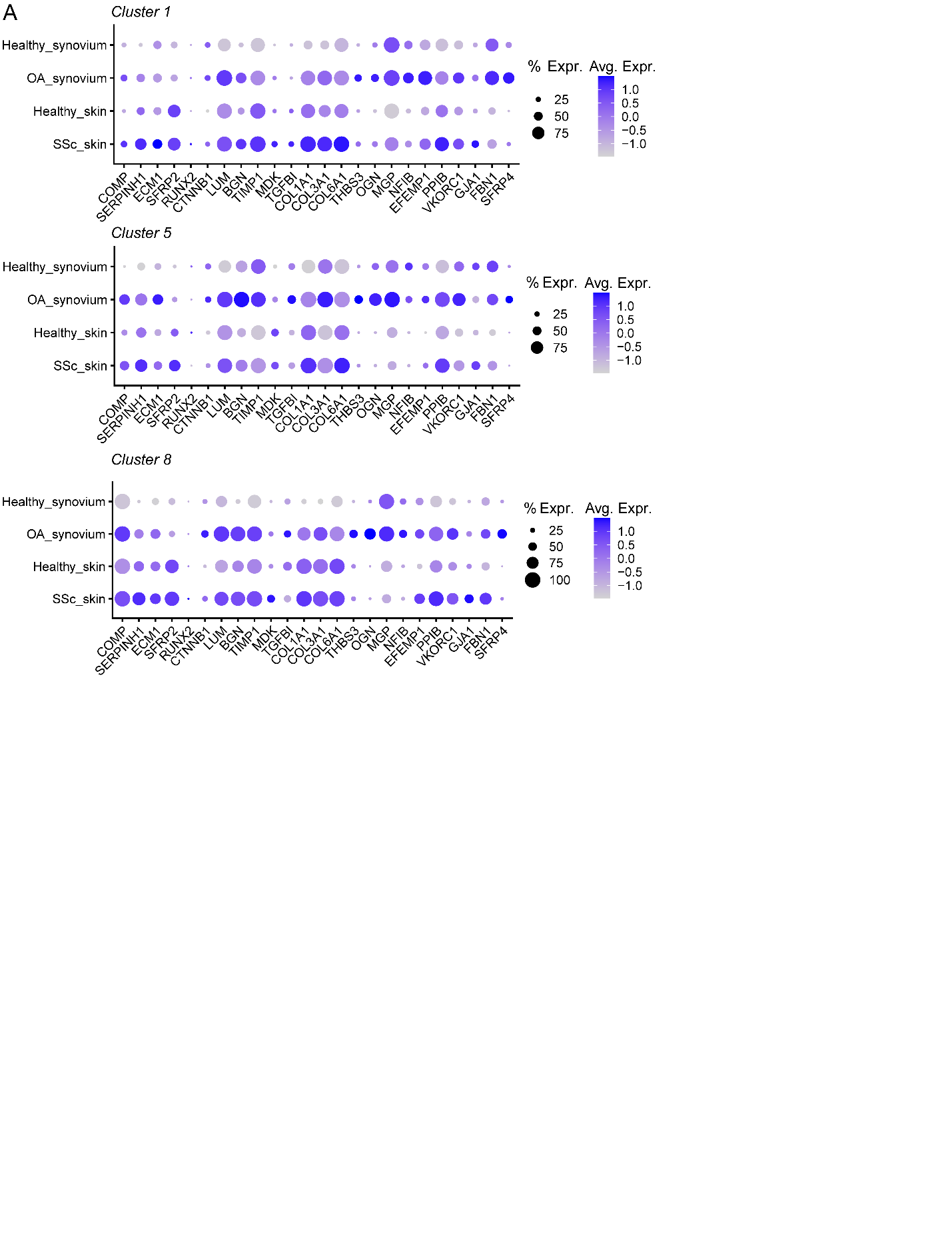

### Supplemental Figure 5: Disease-associated osteochondral gene expression in fibroblast clusters 1, 5, and 8.

**(A)** Dot plots showing expression of osteochondral pathway genes in fibroblast clusters 1 (*MFAP5+*, *PI16+*), 5 (*SFRP2+*, *POSTN+*), and 8 (*SFRP2+*, *COMP+*), stratified by tissue and condition. Genes were derived from the chondrocyte development, cartilage development, and bone development GO:BP terms shown in Figure 4. Dot color indicates scaled average expression, and dot size indicates the percentage of cells expressing each gene.

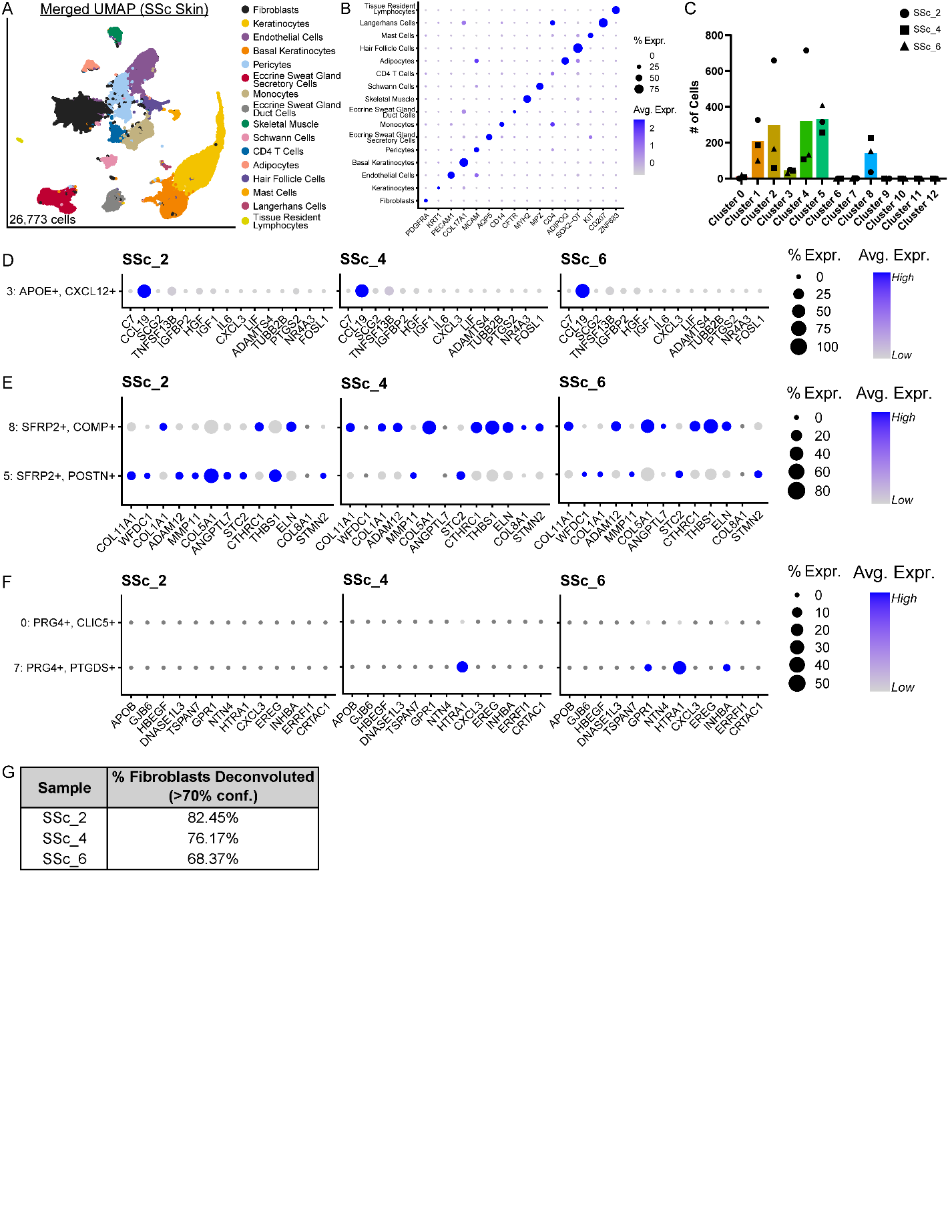

**Supplemental Figure 6: SSc skin Xenium 5K cell-type annotation and fibroblast-subset deconvolution.**

**(A)** UMAP of 26,773 cells merged across three SSc skin Xenium 5K samples, annotated by major cell type. **(B)** Dot plot showing canonical marker-gene expression across major cell types in the merged SSc skin dataset. Dot color indicates scaled average expression, and dot size indicates the percentage of cells expressing each gene. **(C)** Numbers of fibroblasts assigned to each scRNA-seq-derived fibroblast cluster following deconvolution. Symbols represent individual subject samples. **(D–F)** Dot plots showing the expression of selected scRNA-seq cluster-defining genes in deconvoluted fibroblast subsets from each SSc skin sample: **(D)** immune-signaling cluster 3 (*APOE+*, *CXCL12*+); **(E)** myofibroblast-like clusters 5 (*SFRP2*+, *POSTN*+) and 8 (*SFRP2*+, *COMP+*); and **(F)** lining and lining-like clusters 0 (*PRG4*+, *CLIC5*+) and 7 (*PRG4*+, *PTGDS*+). Dot color indicates scaled average expression, and dot size indicates the percentage of cells expressing each gene. **(G)** Percentage of fibroblasts in each SSc skin sample confidently assigned to an scRNA-seq derived fibroblast subset using a posterior probability threshold of ≥ 0.70.

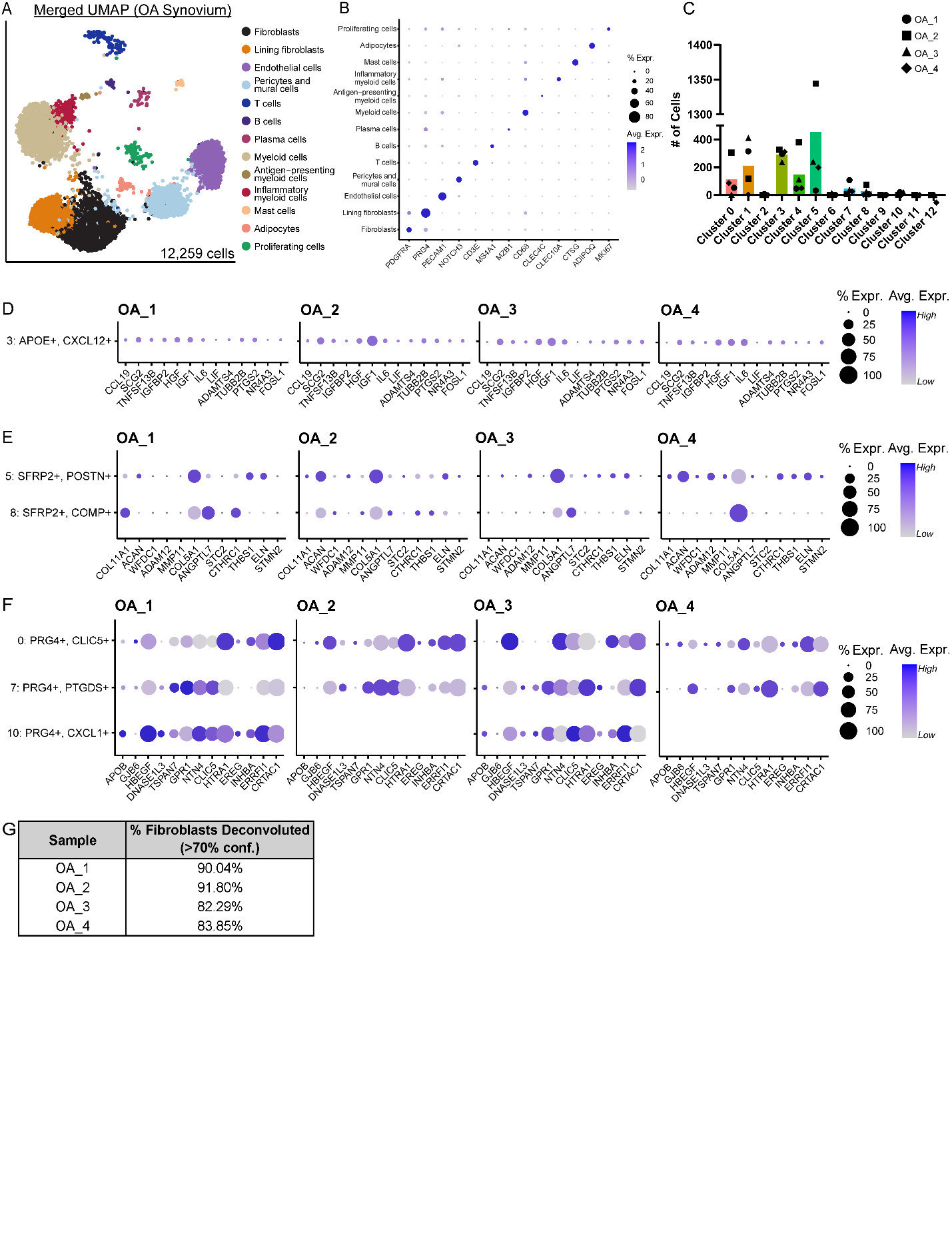

### **Supplemental Figure 7:** OA **synovium** Xenium 5K cell type deconvolution

**(A)** UMAP of 12,259 cells merged across four OA synovium Xenium 5K samples, annotated by major cell type. **(B)** Dot plot showing canonical marker-gene expression used to annotate major cell types in the merged OA synovium dataset. Dot color indicates scaled average expression, and dot size indicates the percentage of cells expressing each gene. **(C)** Numbers of fibroblasts assigned to each scRNA-seq-derived fibroblast cluster following deconvolution. Bars indicate total cell counts, and overlaid symbols represent individual samples. **(D–F)** Dot plots showing expression of selected scRNA-seq cluster-defining genes in deconvoluted fibroblast subsets from each OA synovium sample: **(D)** immune-signaling cluster 3 (*APOE*+, *CXCL12*+); **(E)** myofibroblast-like clusters 5 (*SFRP2*+, *POSTN*+) and 8 (*SFRP2*+, *COMP*+); and **(F)** lining and lining-like clusters 0 (*PRG4*+, *CLIC5*+), 7 (*PRG4*+, *PTGDS*+), and 10 (*PRG4*+, *CXCL1*+). Dot color indicates scaled average expression, and dot size indicates the percentage of cells expressing each gene. **(G)** Percentage of fibroblasts in each OA synovium sample confidently assigned to an scRNA-seq derived fibroblast subset using a posterior probability threshold of ≥ 0.70.

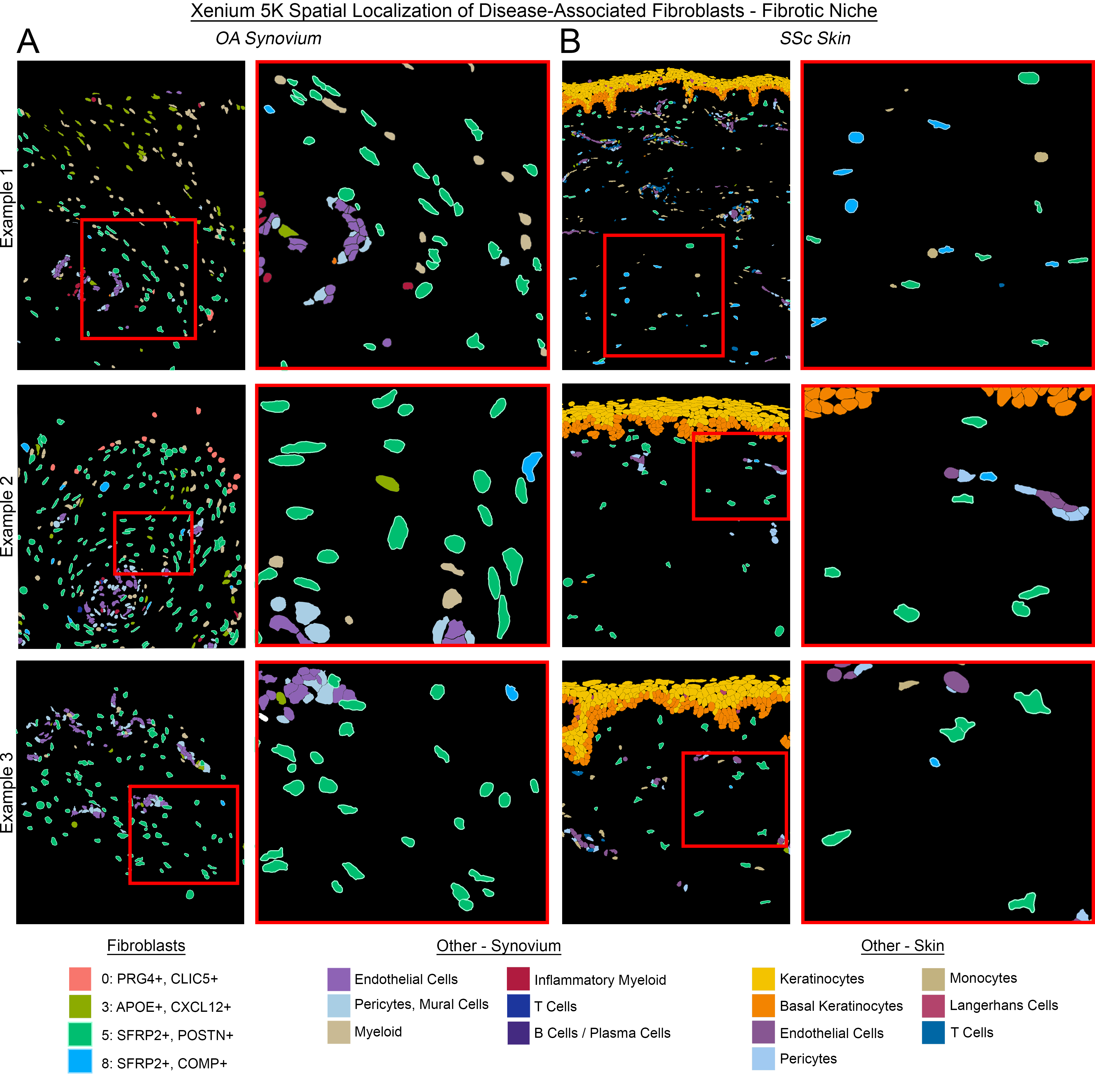

### Supplemental Figure 8: Spatial organization of disease-associated fibroblast subsets within the fibrotic niche in OA synovium and SSc skin

**(A-B)** Additional representative fibrotic regions from Xenium 5K spatial transcriptomic samples of **(A)** OA synovium (Example 1: OA_3; Example 2: OA_2; Example 3: OA_4) and **(B)** SSc skin (Example 1: SSc_6; Example 2: SSc_6; Example 3: SSc_2). Left: Spatial maps showing selected deconvoluted fibroblast subsets and neighboring stromal and immune cell types. Boxes indicate regions enlarged at right. Right: Enlarged views highlighting the distribution of myofibroblast-like clusters 5 (*SFRP2+*, *POSTN+*) and 8 (*SFRP2+*, *COMP+*) within fibrotic tissue regions. Only fibroblasts assigned to a subcluster with a deconvolution probability ≥ 0.70 are shown.

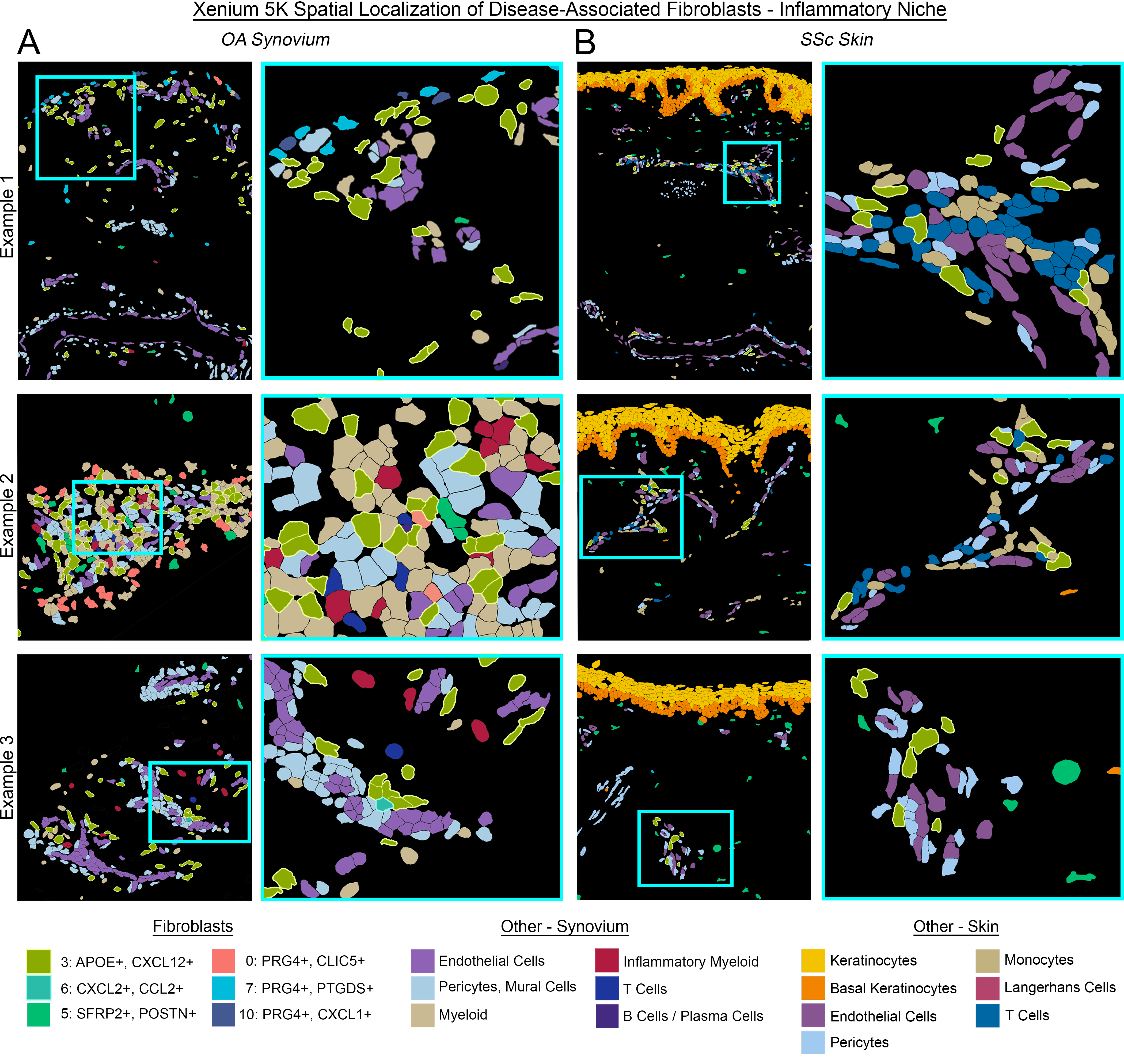

### **Supplemental Figure 9: Spatial organization of disease-associated fibroblast subsets within the inflammatory niche in OA synovium and SSc skin.**

**(A-B)** Additional representative inflammatory regions from Xenium 5K spatial transcriptomic samples of **(A)** OA synovium (Example 1: OA_1; Example 2: OA_4; Example 3: OA_3) and **(B)** SSc skin (Example 1: SSc_2; Example 2: SSc_4; Example 3: SSc_6). Left: Spatial maps showing selected deconvoluted fibroblast subsets and neighboring vascular and immune cell types. Boxes indicate regions enlarged at right. Right: Enlarged views highlighting cluster 3 (*APOE+*, *CXCL12+*) fibroblasts positioned adjacent to endothelial cells, pericytes, and infiltrating immune cells. Only fibroblasts assigned to a subcluster with a deconvolution probability ≥ 0.70 are shown.

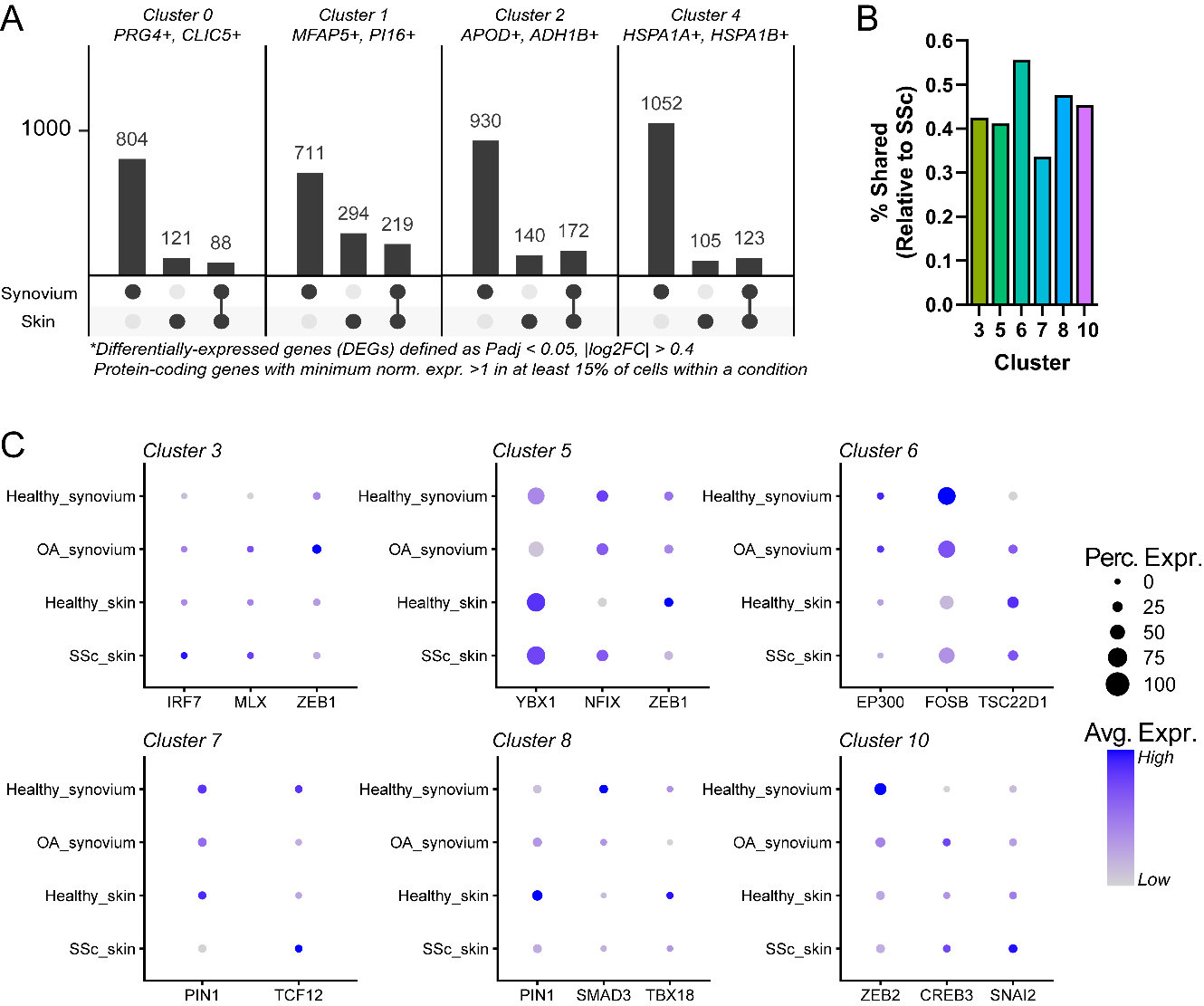

### Supplemental Figure 10: Transcription factor gene expression

**(A)** UpSet plots summarizing pseudobulk differential-expression analyses between diseased and healthy fibroblasts within synovium and skin for fibroblast clusters not included in the disease-associated profibrotic cluster analysis in Figure 6. Bars indicate the numbers of differentially expressed genes (DEGs) unique to OA synovium, unique to SSc skin, or shared between both diseases. DEGs were identified using DESeq2 with a Wald test and *apeglm* log2 fold-change shrinkage (*P_adj_* < 0.05, |log2FC| > 0.4, and expression in >15% of cells in at least one condition). **(B)** Shared DEGs as a percentage of all SSc DEGs for disease-associated fibroblast clusters 3, 5, 6, 7, 8, and 10. **(C)** Expression of the top transcription factors, shown across the corresponding fibroblast clusters and stratified by tissue and condition. TFs with increased predicted activity in ==in both tissues, whereas TFs with decreased predicted activity were required to be expressed in >10% of cells in at least one healthy condition in both tissues. Dot color indicates average expression, and dot size indicates the percentage of cells expressing each TF.
